# A new 1,4-Dihydropyridine-Based L-/T-Type Calcium Channel Inhibitor, HM12, Provides in Vivo and in silico Cardioprotective Effects in Doxorubicin-Treated Rats

**DOI:** 10.1101/2025.09.21.677599

**Authors:** Oluwafemi Ezekiel Kale, Miyase Gözde Gündüz, Ifabunmi Oduyemi Osonuga, Olufunsho Awodele, Martins Ekor

**Author notes:** **Correspondence:** Oluwafemi Ezekiel Kale, Department of Pharmacology, Olabisi Onabanjo University, Obafemi Awolowo College of Health Sciences, Sagamu Campus, P.O.Box 2001, Ago-Iwoye, Ogun State, Nigeria.

## Abstract

**Background:** Doxorubicin (DOX), an anthracycline anticancer agent, has limited use due to its cardiotoxicity via oxidative stress and mitochondrial dysfunction. This study evaluated HM12, a 1,4-dihydropyridine calcium channel blocker (CCB) derivative, for protective effects against DOX-induced toxicity.

**Methods:** Eight groups of adult male Wistar rats received saline, DOX (20 mg/kg), HM12 (5 or 20 mg/kg), nifedipine (NFD, 20 mg/kg), or combinations of DOX with HM12 or NFD. Assessments included blood biochemistry, cardiac biomarkers, oxidative-antioxidant indices, renal and hepatic function tests, and organ histology. In silico docking was performed using human topoisomerase IIβ (3QX3).

**Results:** DOX induced marked cardiotoxicity, evidenced by elevated TNF-α, IL-6, C-RP, LDH, and cardiac MDA. Renal and hepatic toxicity were also observed, with increased MDA levels. HM12 improved heart weight and significantly reduced IL-6, C-RP, and LDH, though not TNF- α. Antioxidant defenses improved, with increased glutathione, catalase, and superoxide dismutase activity. HM12 offered limited protection to renal and hepatic tissues. Histologically, the higher HM12 dose ameliorated cardiac damage. Notably, in silico HM12 exhibited greater binding affinity than NFD and engaged in distinct interaction patterns with 3QX3 that were not observed with DOX.

**Conclusion:** HM12 shows cardioprotective effects against DOX-induced toxicity, likely via antioxidant enhancement and modulation of inflammatory markers, though its protection of renal and hepatic tissues is limited.

## 1 INTRODUCTION

Doxorubicin (DOX), also known as adriamycin, is an anthracycline antibiotic derived from *Streptomyces peucetius*. It is widely recognized for its efficacy against solid tumors and its use in the chemotherapeutic management of breast carcinomas (Sallustio & Boddy, 2021; Bisht et al., 2025). However, the clinical application of DOX is significantly limited by its dose-dependent and irreversible cardiotoxicity, which can progress to cardiomyopathy and congestive heart failure (Saleh et al., 2021; Sheibani et al., 2022). Approximately 10% of cancer survivors treated with DOX or its derivatives develop cardiovascular complications either during or after chemotherapy (Gil-Gil et al., 2021).

While DOX’s principal adverse effects manifest as systemic toxicity (Kamińska & Cudnoch-Jędrzejewska, 2023), its deleterious impact particularly on the heart and other vital organs even at therapeutic doses, highlights the urgency of addressing its cardiotoxic potential (Asnani, 2021). The mechanisms underlying DOX-induced cardiotoxicity are complex and multifactorial (Rawat et al., 2021). Studies have identified a cascade of biological events influenced by various factors, often overlapping with other cellular processes. Notably, DOX has been shown to deplete myocardial antioxidant enzymes and cause genetic alterations (Rawat et al., 2021; Kong et al., 2022). This toxicity is largely mediated by the generation of highly reactive oxygen species (ROS), which bind covalently to vital biomolecules (Zhang et al., 2021; Kong et al., 2022). These ROS induce damaging complexes of proteins, lipids, and DNA, weakening endogenous antioxidant defenses and leading to cellular injury (Antonucci et al., 2021).

Despite being toxic to both cancerous and normal cells, the precise mechanism of DOX-induced cell death remains unclear. Among the proposed mechanisms, the dysregulation of calcium homeostasis has emerged as a central factor (Awad et al., 2021; Zhang et al., 2021; Shinlapawittayatorn et al., 2022; Belger et al., 2024). DOX is believed to interfere with calcium signaling, triggering abnormal excitability in tumor cells and intracellular organelles (Santostasi et al., 1991; Zhang et al., 2021; Bhatt et al., 2025). Further evidence suggests that DOX forms adducts with calcium-calmodulin complexes, disrupting their function and driving cellular dysfunction (Maleki et al., 2022; Morciano et al., 2022; Shinlapawittayatorn et al., 2022; Belger et al., 2024). Calcium-related mechanisms have also been implicated in DNA damage, mitochondrial dysfunction, inflammation, sympathetic overstimulation, immunogenic responses, and apoptosis (Santostasi et al., 1991; Mordente et al., 2012; Kong et al., 2022). Collectively, these processes contribute to early cessation of chemotherapy due to toxicity (Yuan et al., 2021).

At present, only a limited number of studies have identified agents capable of mitigating DOX-induced cardiotoxicity without compromising its anticancer efficacy (Abushouk et al., 2017; Awad et al., 2021; Saleh et al., 2022). To date, no drug has been conclusively shown to prevent this toxicity, largely due to incomplete understanding of its underlying mechanisms. Among the most promising candidates are the calcium channel blockers (CCBs), a diverse class of drugs widely used to manage cardiovascular diseases and hepato-renal conditions (Hassan et al., 2020; Karthick et al., 2022).

The regulation of calcium homeostasis is a complex process involving numerous signaling pathways (Huber et al., 2021). Although the precise role of calcium ion channels in DOX toxicity remains to be fully elucidated, various studies suggest that CCBs may confer protective effects against DOX-induced injury (Ikeda et al., 2019; Panneerpandian et al., 2021; Shinlapawittayatorn et al., 2022; Sripusanapan et al., 2025). Some reports further propose that the mobilization of calcium ions either directly or indirectly is essential for mitigating the oxidative stress induced by DOX (Minotti et al., 2004). While expert reviews continue to update strategies for managing both acute and chronic DOX toxicities (Kong et al., 2022; Aloss & Hamar, 2023), no ideal adjuvant has yet been found. Nonetheless, numerous preclinical and clinical studies have identified a variety of candidate agents (Aloss & Hamar, 2023).

Hemodynamic studies have highlighted the ability of 1,4-dihydropyridines (DHPs), a major class of CCBs to reduce total arterial resistance and enhance cardiac performance (Bertram-Ralph & Amare, 2021). Studies have identified and reported various potential agents capable of modulating doxorubicin (DOX)-induced cardiotoxicity without reducing its antitumor efficacy. Among the most promising and extensively studied clinical drugs are CCBs, a significant class of medications used in the treatment of a range of cardiovascular diseases, as well as in modulating hepato-renal functions. Currently, no ideal adjuvant exists for managing DOX-induced toxicity. But, CCBs have demonstrated antioxidant properties in both in vitro and in vivo models (Kale et al., 2017; Heravi & Zadsirjan, 2022; Jones et al., 2024). In one study, a DHP-based compound, HM12, was shown to effectively block both L- and T-type calcium channels (Aygün et al., 2019).

Additionally, DOX actions similar to other etoposide-class of drugs, to inhibit the Topoisomerase II isoenzymes, Top2α and Top2β, in mammalian cells have been updated (Linders et al., 2024; Bhutani et al., 2025). These isoenzymes are regulated by distinct cellular mechanisms. Because Top2β which regulatory role in postmitotic cells is expressed in adult cardiac tissue, it has been implicated in the cardiotoxic side effects of DOX (Vejpongsa & Yeh, 2014; Bhutani et al., 2025). In contrast, Top2α is typically upregulated in proliferating tumor cells (Bhutani et al., 2025). DOX is known to stabilize the Top2β-DNA cleavage complex, leading to early apoptosis and consequently cardiac problems (Vejpongsa & Yeh, 2014; Linders et al., 2024).

Currently, there is limited information on the relationship between dihydropyridines and DOX-induced cardiotoxicity through effects on the Top2β isoenzyme. Although dihydropyridine calcium channel blockers (CCBs) have been reported to confer cardioprotective benefits (Bertram-Ralph & Amare, 2021; Jones et al., 2024), some evidence suggests they may negatively affect Top2β and potentially contribute to cardiotoxicity and altered gene regulation (Ganapathi & Ganapathi, 2013; Delint-Ramirez et al. 2022;). On the other hand, HM12, a compound with both L- and T-type calcium channel blocking activity may offer improved *in* vitro properties over conventional dihydropyridines.

Therefore, in furtherance of this observation, we sought to evaluate the potential chemopreventive role of HM12 against DOX-associated cardiotoxicity. Also, we hypothesize that while conventional dihydropyridine CCBs may share cytotoxic effects on Top2β similar to those of etoposides, HM12 may differ in its interaction. To investigate this, we first employed in silico molecular fingerprinting to assess whether dihydropyridine CCBs and DOX may potentiate cardiotoxic effects via human Top2β. Further, we evaluated the potential interaction of HM12 with human Top2β. This is to ascertain it effect, elucidate its modulatory mechanisms and enhance its clinical utility.

## 2 MATERIALS AND METHODS

### 2.1 Drugs and chemicals

Doxorubicin was purchased from Neon Laboratories Limited (28 Mahal Industrial Estate, M. Caves Road, Andheri (East), Mumbai 400093, India). Nifedipine was obtained from Unicure Remedies Pvt. Ltd. (India). The compound HM12 was synthesized at the Department of Pharmaceutical Chemistry, Faculty of Pharmacy, Hacettepe University, Ankara, Turkey. Thiobarbituric acid (TBA) and 5,5′-dithiobis-(2-nitrobenzoic acid) (Ellman’s reagent) were procured from Sigma-Aldrich Chemical Co. (St. Louis, MO, USA). Reduced glutathione (GSH), metaphosphoric acid, and trichloroacetic acid (TCA) were purchased from J.T. Baker (Phillipsburg, NJ, USA). Tumor necrosis factor-alpha (TNF-α, Rat) ELISA kit (45- TNFRTE01.1) was obtained from Eagle Biosciences inc. (USA). The Rat C-reactive protein (C- RP) ELISA kit (SEA821Ra) was obtained from Katy, TX 77494 (USA). Lactate dehydrogenase (LDH) reagent was obtained from Oroboros Instruments Corp. (Schöpfstr. 18, A-6020 Innsbruck, Austria; MiPNet08.18). The Rat Interleukin-6 (IL-6) ELISA kit (IL621-K01) was purchased from The Eagle Biosciences (20A Northwest Blvd., Suite 112, Nashua, NH 03063, USA). All other chemicals and reagents used were of analytical grade.

### 2.2 Synthesis

The synthesis of HM12 (2-[(2-methylacryloyl)oxy]ethyl 4-(3,5-dibromo-2-hydroxyphenyl)- 2,6,6-trimethyl-5-oxo-1,4,5,6,7,8-hexahydroquinoline-3-carboxylate) was previously described by Aygün Cevher et al. (2019). In brief, equimolar amounts of 4,4-dimethyl-1,3- cyclohexanedione, 3,5-dibromo-2-hydroxybenzaldehyde, and 2-(methacryloyloxy)ethyl acetoacetate, along with an excess amount of ammonium acetate, were heated in ethanol under microwave irradiation (Figure 1). Upon completion, the reaction mixture was poured into ice water. The resulting precipitate was purified by recrystallization from an ethanol–water mixture to yield HM12.

**Figure 1:**
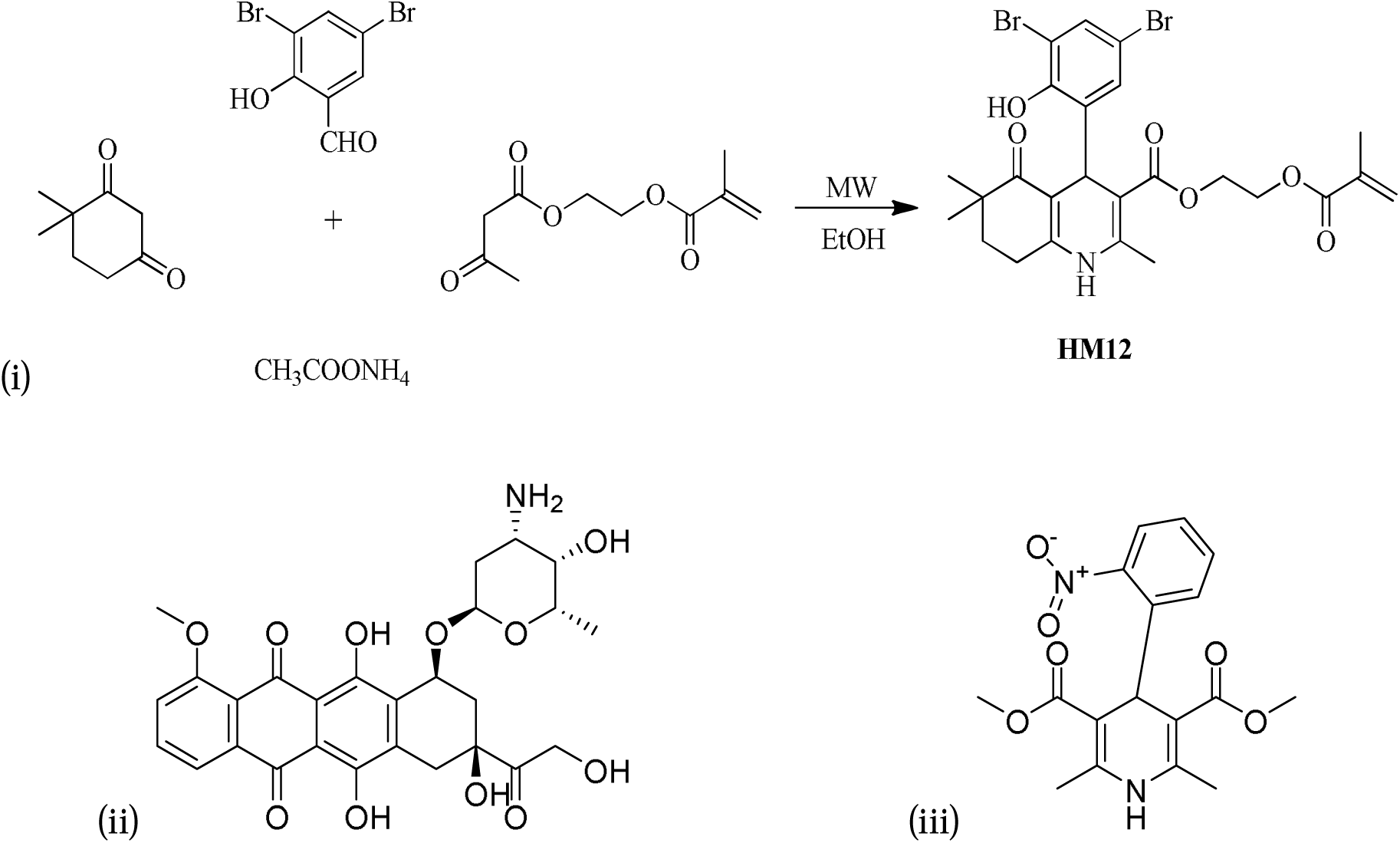
Chemical structures (i) The synthetic route to HM12 (ii) Doxorubicin (iii) Nifedipine.

### 2.3 Animals

Male Wistar albino rats were procured from the Animal House of the Faculty of Basic Medical aand Clinical Sciences, Babcock University, Nigeria. The animals were housed under standard laboratory conditions of ambient temperature and humidity, with a 12-hour light-dark cycle within the experimental animal-handling facility of the Department of Pharmacology,

Therapeutics and Toxicology, College of Health Sciences, Olabisi Onabanjo University. Throughout the acclimatization period and the duration of the experiment, the rats were provided with a pelleted rat diet (Ladoke Akintola Grower Mash, Nigeria) and water *ad* libitum. This study adhered to the established guidelines for the care and use of laboratory animals in biomedical research and education, as approved by the Institute of Laboratory Animal Resources, National Research Council (DHHS Publication No. NIH 86-23, 1985).

### 2.4 Experimental design and treatment

Forty-eight (48) male rats (weighing 120 - 150 g) were randomly divided into eight groups, with six rats per group. Doxorubicin (DOX) toxicity was induced using a subacute administration protocol, as described by Iqbal et al. (2008). Rats in Groups 6, 7, and 8 were pretreated with the test compound one hour prior to DOX administration. All treatments were administered orally once daily for seven consecutive days. The test compound, HM12, was dosed based on human equivalent dosing for a 70 kg adult male, at both lower (5 mg/kg/day) and upper (20 mg/kg/day) therapeutic limits. On day 8, after an overnight fast, the rats were weighed and blood samples were collected via the retro-orbital plexus using capillary tubes and transferred into lithium heparin-coated tubes for plasma separation or into EDTA tubes to prevent coagulation. Animals were euthanized via cervical dislocation following ketamine anaesthesia (10%). Whole samples were centrifuged at 4000 rpm for 10 minutes at room temperature. Biochemical and histological assessments were subsequently performed. Treatment was as follow:

Group 1: Control normal saline (1 ml/kg/day);

Group 2: Doxorubicin (DOX, 20 mg/kg/day, i.p.) positive control rats received a single cardiotoxic dose of DOX.

Group 3: HM12 upper dose (UDHM12, 20 mg/kg/day, p.o.), rats was administered an upper dose HM12.

Group 4: HM12 lower dose (LDHM12, 5 mg/kg/day, p.o.), rats were treated with low dose HM12.

Group 5: Nifedipine (NFD, 20 mg/kg/day, p.o.), rats were treated with NFD (20 mg/kg, p.o.).

Group 6: LDHM12 (5 mg/kg/day) + DOX (20 mg/kg/day, i.p.);

Group 7: UDHM12 (20 mg/kg/day) + DOX (20 mg/kg/day, i.p.); Group 8: NFD (20 mg/kg/day) + DOX (20 mg/kg/day, i.p.);

### 2.5 Ethical concerns in animal study

All animal experiments and protocols were conducted in accordance with the guidelines of the National Institutes of Health (NIH, 2000) for the care and use of laboratory animals. All procedures adhered to the protocols approved by the Nigerian Institutional Animal Care and Use Committee. The animals were certified fit for experimentation by the Institution’s Animal Health Officers, following the guidelines outlined by Kilkenny et al. (2010) for reporting animal research. Institutional procedures of the Faculty of Basic Medical Sciences, College of Health Sciences, Olabisi Onabanjo University, Ogun State, Nigeria, were strictly followed. Ethical approval was obtained from the University Health Research Ethics Committee (BUHREC 376/18) prior to 1,4-Dihydropyridine-Based L-/T-Type Calcium Channel Inhibitor research study. After the experiments, all animal carcasses were disposed of by burial at least two feet below the natural ground surface, covered with lime and disinfectant, and then sealed with soil.

### 2.6 Binding Affinity and molecular docking of HM12 with human TOP2B

The Human topoisomerase II beta in complex with DNA and etoposide (3QX3), known for its isomerase inhibition, is from the Protein Data Bank. Protein (3QX3) structure was visualized, purified and analyzed using PyMOL 2.6.0 and Biovia Discovery Studio 2024. 3QX3 chain A was analysed and prepared for Autodock process. Chemical structures of doxorubicin and nifedipine compounds were retrieved from PubChem and converted to PDB format using Open Babel. HM12 was drawn and made MDL SDF file using ChemDraw and MarvinSketch. HM12 smile was retrived from EMBL-EBL website (Wellcome Genome Campus, Hixton Cambridgeshire CB10 1SK, UK). These PDB files, with Gasteiger charges, were converted to PDBQT format for protein-ligand docking using AutoDock Vina v4.2.6. PyRx virtal screening tool was used to minimize and obtained pdbqt format of all ligands and make macromolecule of protein. Docking results were visualized with PyMOL 2.6.0, and interaction analyses were performed in Biovia Discovery Studio to generate 2D diagrams and 3D visualizations.

### 2.7 Statistics

Statistical analysis was performed using SPSS (version 20) and GraphPad Prism (version 6). Data are presented as mean ± standard error of the mean (SEM). One-way analysis of variance (ANOVA) was used to compare groups, followed by Tukey’s post hoc test for multiple comparisons. A p-value of < 0.05 was considered statistically significant.

## 3 RESULTS

### 3.1 Heart Weight

The heart weight of the rats was expressed relative to their body weight (Fig. 2). Doxorubicin (DOX) intoxication significantly increased (p < 0.01) heart weight in the control group. Treatment with UDHM12 reduced this elevation by 34.3% compared to the DOX control group. However, neither HM12 nor NFD treatment was able to reverse the loss in body weight observed in the treated animals (Fig. 3).

**Figure 2.**
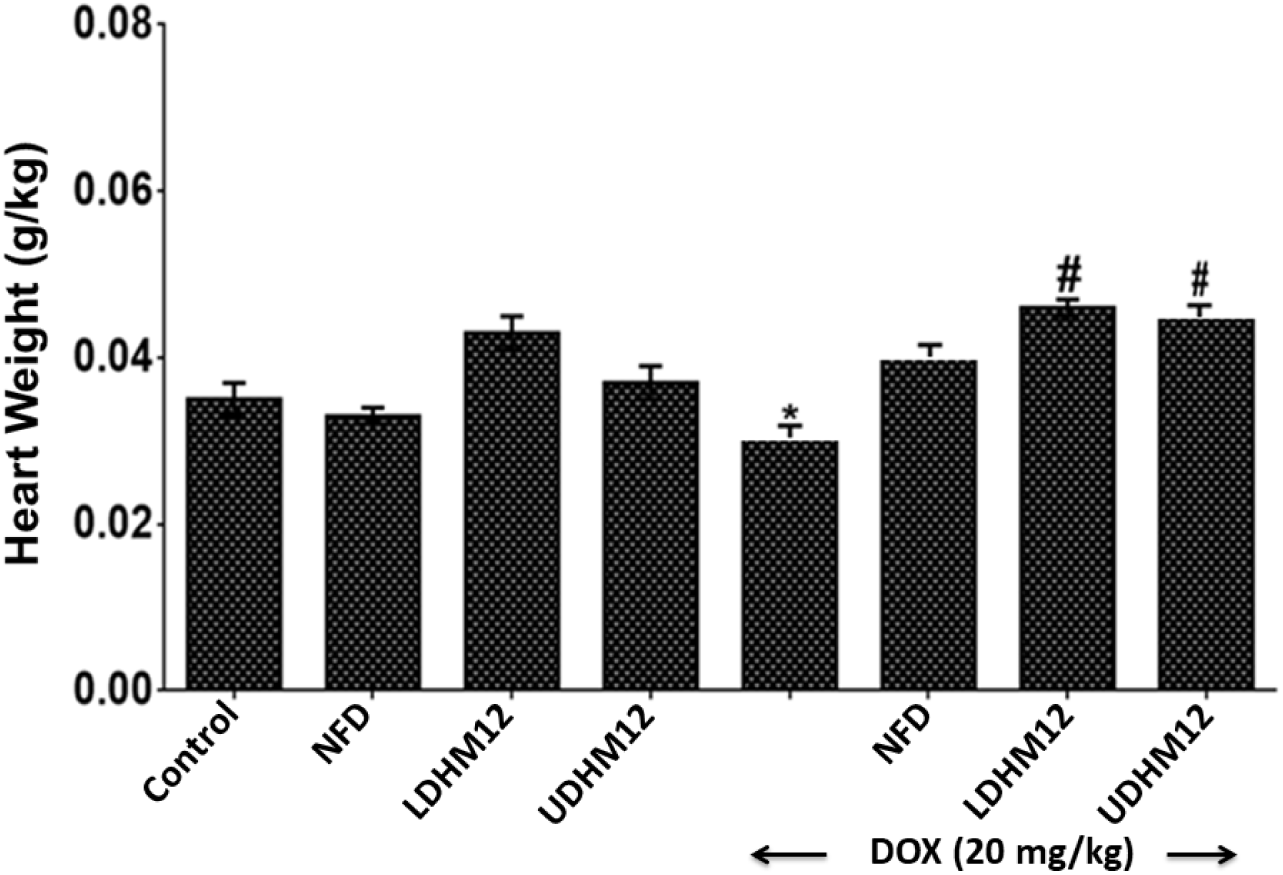
Effect of HM12 on Heart Weight to Body Weight Ratio in Normal and Doxorubicin (DOX)-Treated Male Rats. Results are expressed as Mean ± Standard Error of the Mean (SEM), with n = 6 per group. The control group received saline water (1 ml/kg). *p < 0.01 versus control (saline) group; ^#^p < 0.05 vs. DOX control group.

**Figure 3.**
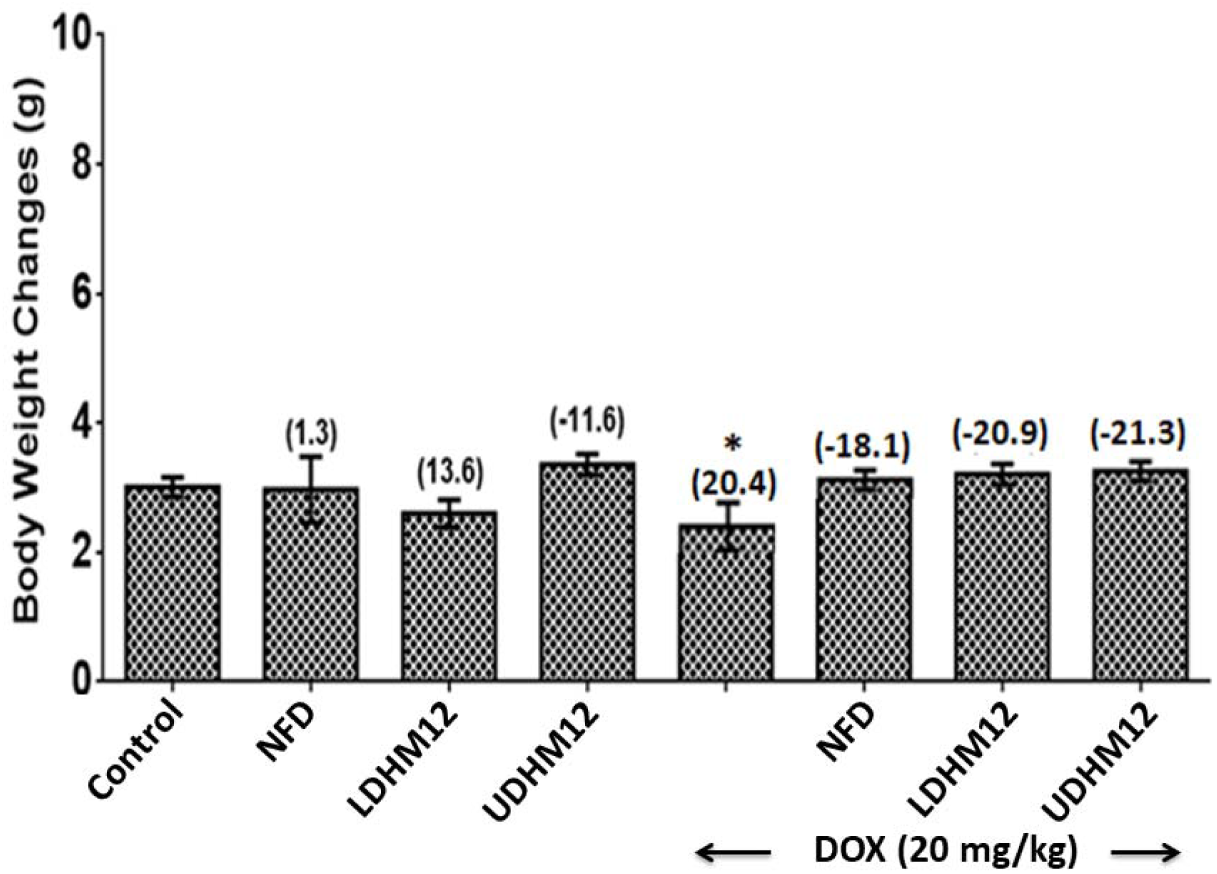
Effect of HM12 on Body Weight Changes in Normal and Doxorubicin (DOX)- Treated Male Rats. Data are presented as Mean ± Standard Error of the Mean (SEM), with n = 6 animals per group. The Δ indicates the change in body weight (Final - Initial). Control animals received saline water (1 ml/kg). *\**p < 0.01 versus control (saline) group; ^#^*p* < 0.05 vs. DOX- treated control group. Values in parentheses represent percentage changes relative to the control (saline) group.

### 3.2 C-RP and LDH levels

The effect of HM12 on plasma C-RP levels is shown in Fig. 4. The DOX-only group exhibited significantly elevated C-RP levels (p < 0.05), with a 46% increase compared to the control (normal saline) group. In contrast, pretreatments with LDHM12 or UDHM12 before DOX intervention resulted in lowered C-RP levels compared to the control groups. The effect of HM12 on plasma LDH levels is shown in Fig. 5. DOX treatment significantly increased LDH levels by 74.7% (p < 0.05) compared to the normal saline control group. Both lower and the higher doses of HM12 effectively reduced the elevated LDH levels in treated rats compared to the DOX control group.

**Figure 4:**
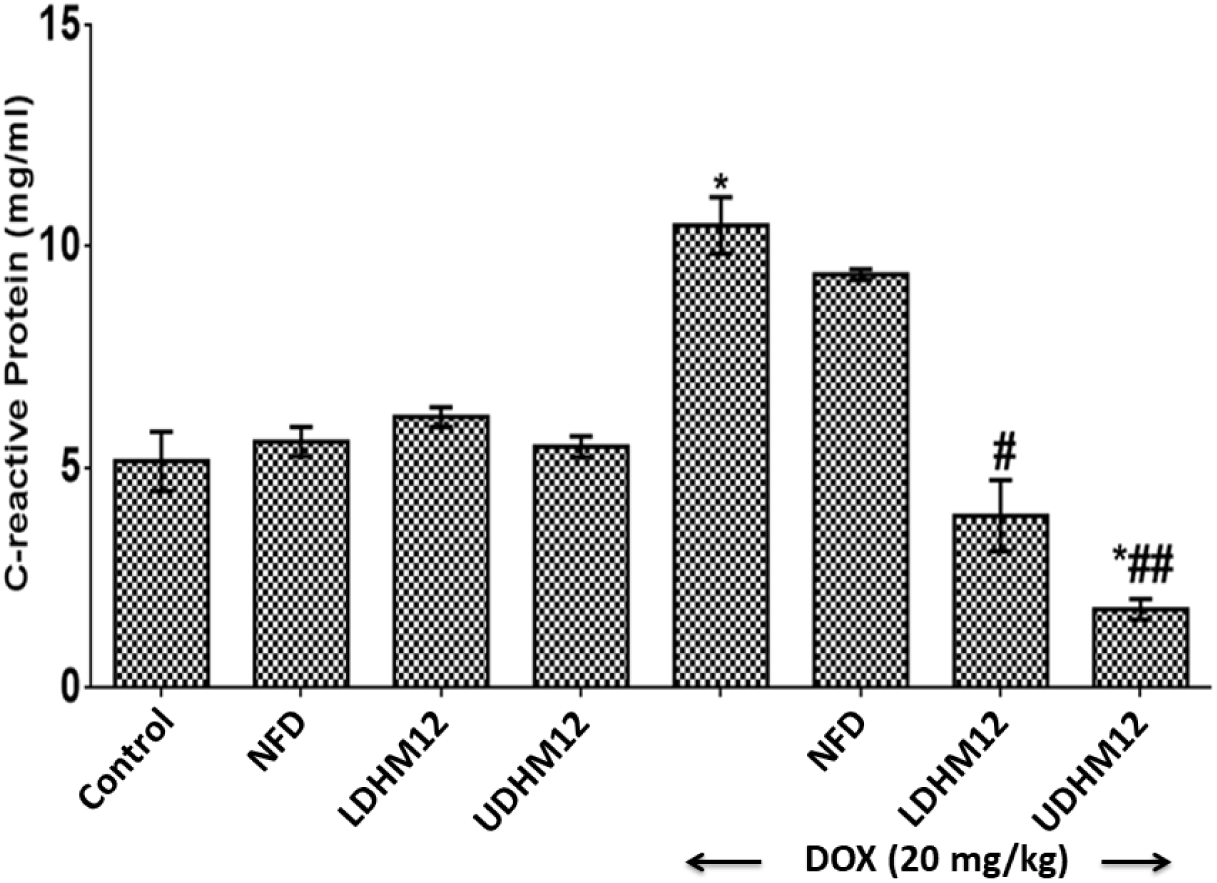
Effect of HM12 on Plasma C-Reactive Protein (C-RP) Levels in Normal and Doxorubicin (DOX)-Treated Male Rats. Data are expressed as mean ± standard error of the mean (SEM), n = 6 per group. Control group received normal saline (1 ml/kg). Plasma C-RP concentrations are presented in ng/ml. \**p* < 0.05 compared with control (saline) group. ^#^*p* < 0.05 compared with control (saline) group. ^##^*p* < 0.05 compared with DOX-treated group.

**Figure 5:**
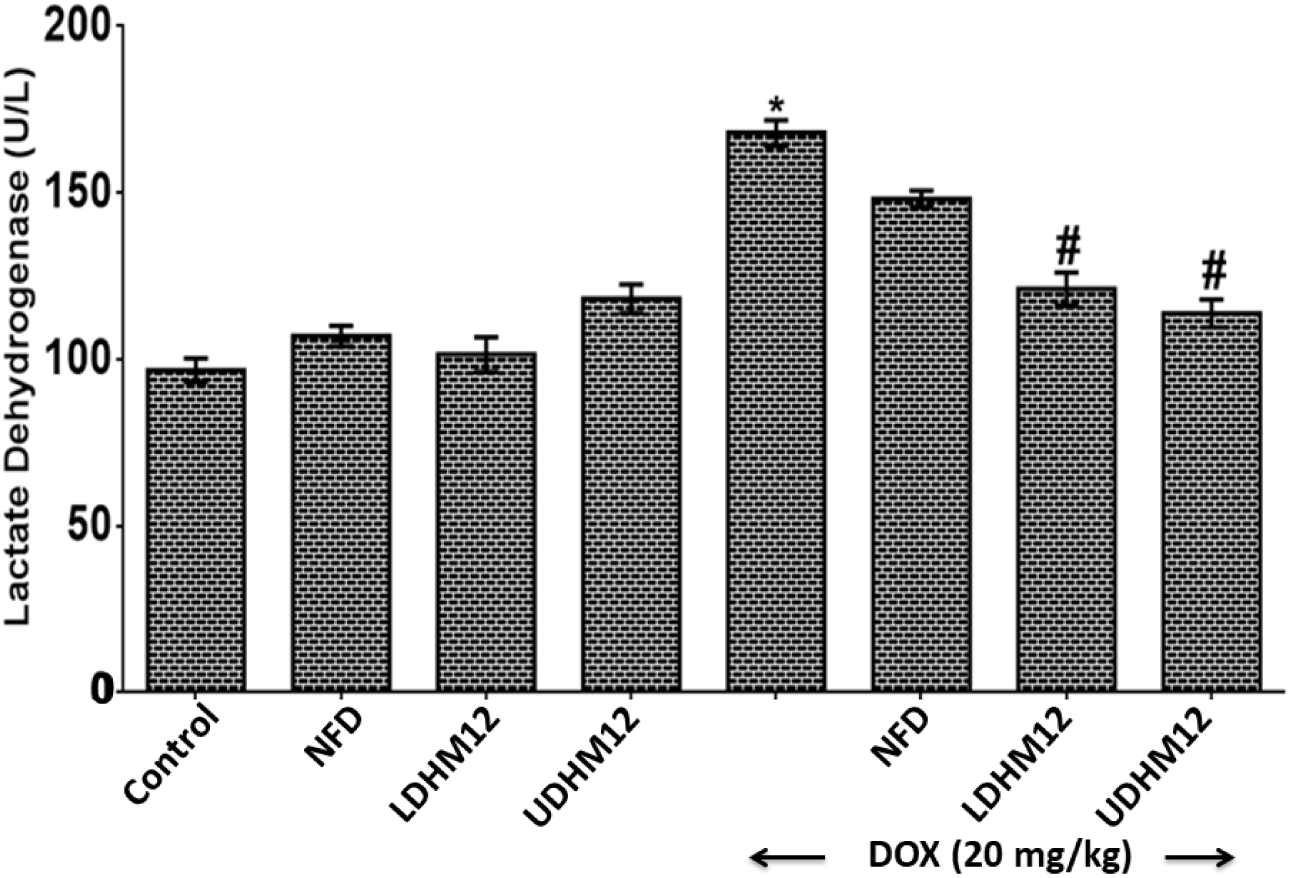
Effect of HM12 on Plasma Lactate Dehydrogenase (LDH) Levels in Normal and Doxorubicin (DOX)-Treated Male Rats. Data are presented as Mean ± Standard Error of Mean (SEM), n = 6 per group. Control group received normal saline (1 ml/kg). LDH activity is expressed in units per liter (U/l). \**p* < 0.05 versus control (saline) group. ^#^*p* < 0.05 versus DOX-treated control group.

**Figure 6:**
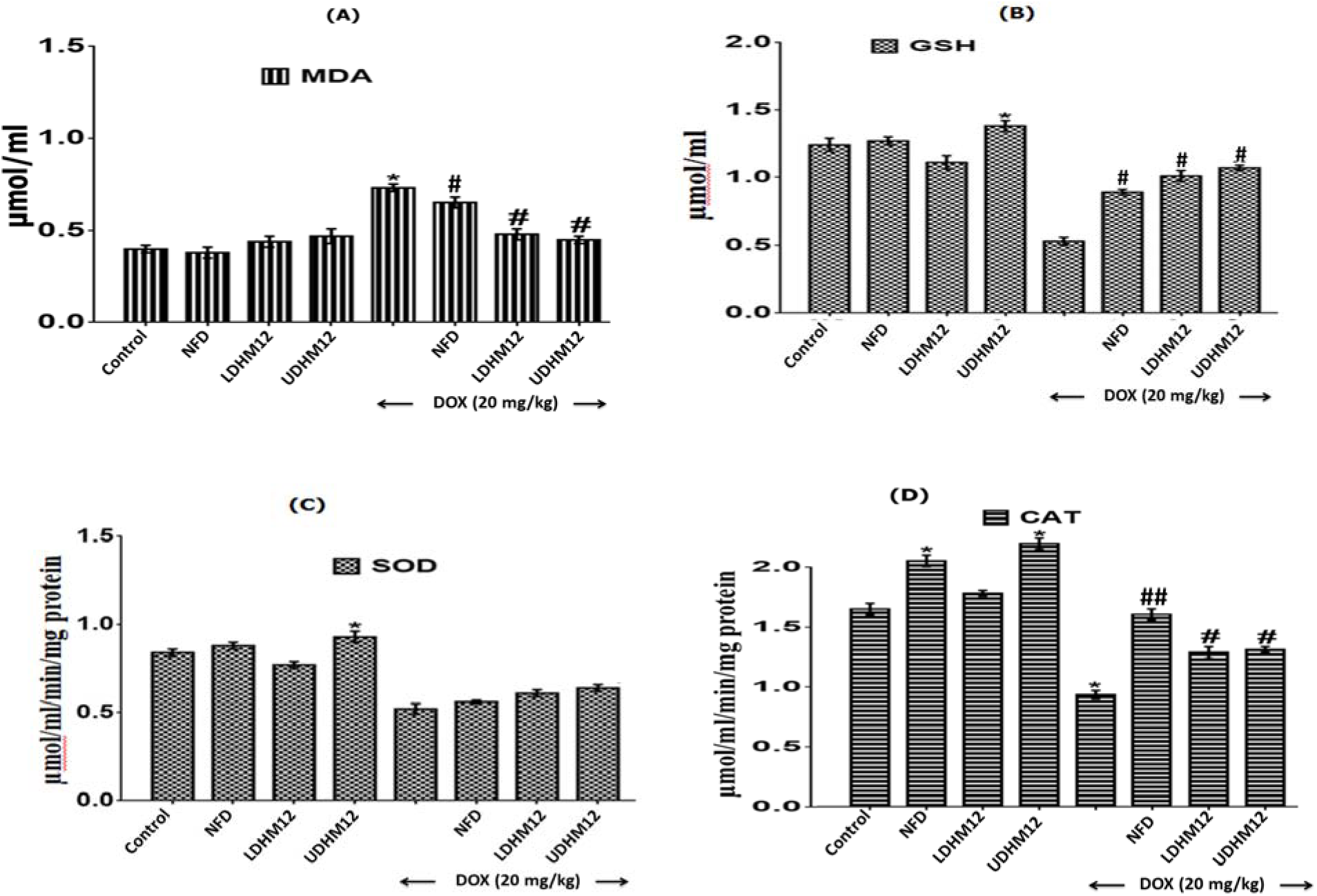
Effect of HM12 on Cardiac Oxidative Stress Markers in Normal and Doxorubicin (DOX)-Treated Male Rats. The effects of HM12 treatment on (a) lipid peroxidation (malondialdehyde, MDA), (b) reduced glutathione (GSH) levels, (c) superoxide dismutase (SOD) activity, and (d) catalase (CAT) activity in cardiac tissues were evaluated in both normal and DOX-treated male rats. Results are expressed as Mean ± Standard Error of Mean (SEM), with n = 6 animals per group. Control group received normal saline (1 ml/kg). MDA: lipid peroxidation, GSH: Reduced glutathione, SOD: Superoxide dismutase and CAT: Catalase. \**p* < 0.05 compared to the saline water control group. ^#^*p* < 0.05 compared to the DOX control group.

### 3.3 Cardiac Lipid Peroxidation Levels and Antioxidants Parameters

The DOX-untreated group showed a significant increase of 78.1% in cardiac lipid peroxidation levels, as measured by malondialdehyde (MDA), compared to the normal control group. Treatment with HM12 significantly reduced MDA levels by 34.2% and 38.4%, respectively, compared to the DOX control group. Additionally, DOX induction caused a marked decrease in cardiac antioxidant defenses: glutathione (GSH) levels dropped by 57.3%, catalase (CAT) activity decreased by 43.3%, and superoxide dismutase (SOD) activity declined by 39.5%. However, treatment with HM12 significantly improved cardiac GSH and CAT levels by at least 50% similar to NFD, a clinical agent, when compared to the DOX control group.

### 3.4 TNF-**α** levels

The effect of HM12 on plasma TNF-α levels in DOX-induced rats is illustrated in Fig. 7. Treatment with DOX caused a significant increase (35.7%) in TNF-α levels compared to the control saline water group. Administration of HM12 at low dose (LDHM12 + DOX) dose did not result in any improvement in TNF-α levels. In contrast, the UDHM12 + DOX or NFD + DOX treatment significantly reduced (p < 0.05) TNF-α levels compared to the DOX-only control group.

**Figure 7:**
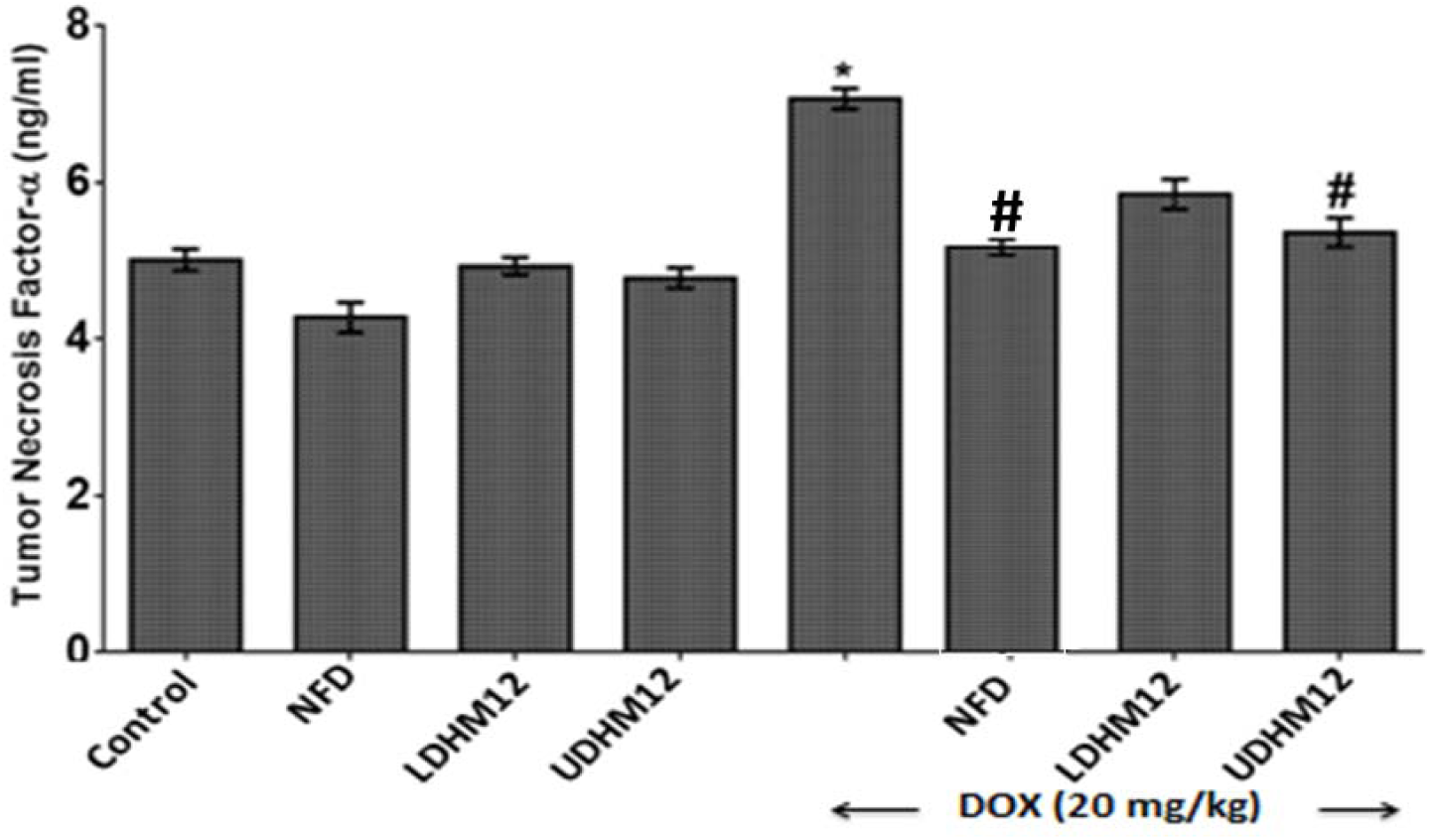
Effect of HM12 on Plasma Tumor Necrosis Factor-Alpha (TNF-α) Levels in Normal and Doxorubicin (DOX)-Treated Male Rats. Plasma levels of TNF-α (ng/ml) were measured in normal and DOX-treated male rats and are presented as Mean ± Standard Error of Mean (SEM), with n = 6 per group. The control group received saline (1 mL/kg). *p < 0.05 versus saline water control, ^#^p < 0.001 versus untreated DOX group.

### 3.5 IL-6 levels

The change in plasma IL-6 levels following administration of HM12 in DOX-induced rats is shown in Fig. 8. DOX treatment caused a significant 36% increase in IL-6 levels compared to the control saline group. However, rats treated with UDHM12 + DOX and NFD + DOX exhibited significantly reduced IL-6 levels up to 30% each compared to the DOX-only group.

**Figure 8:**
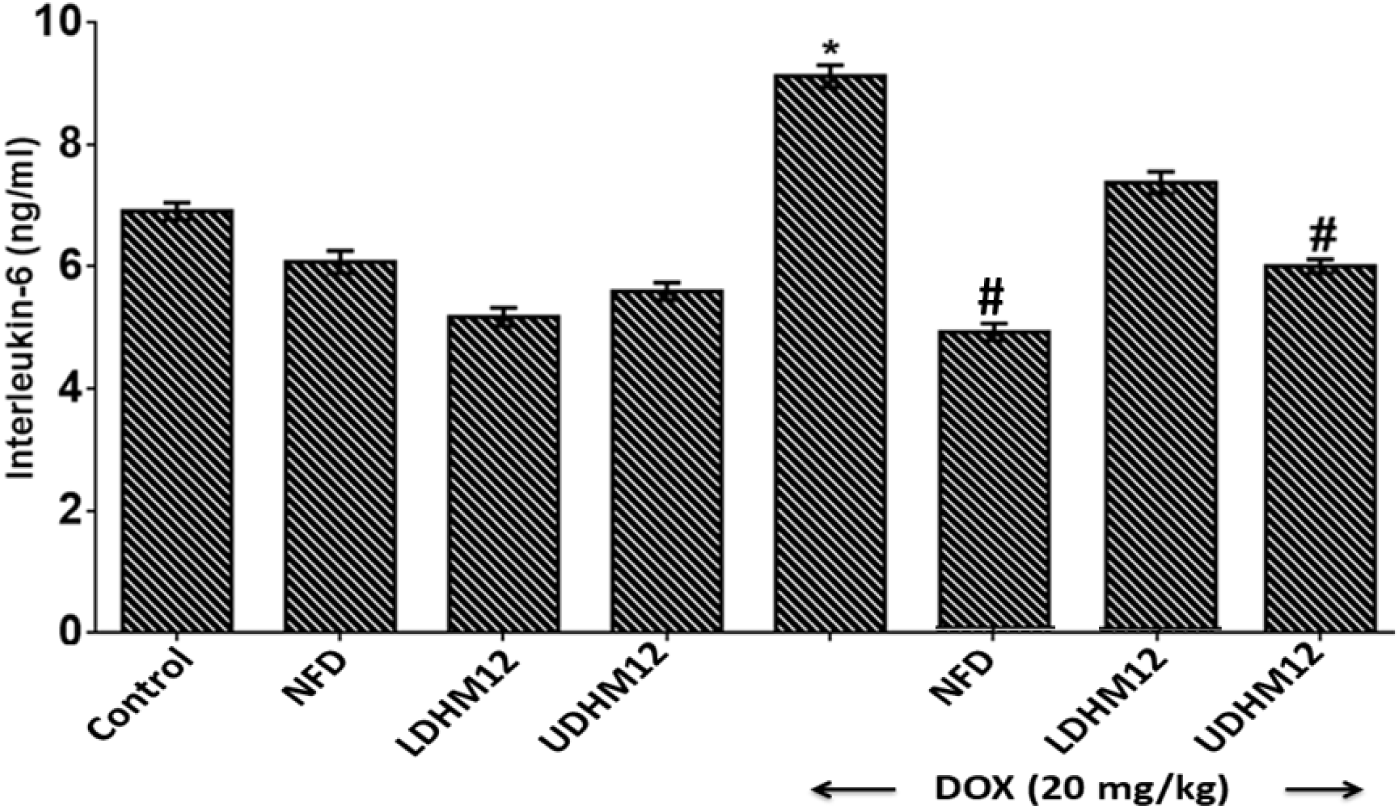
Effect of HM12 on plasma interleukin (IL-6) in normal and doxorubicin (DOX)-treated male rats. Results represented as Mean ± Standard Error of Mean (SEM). n = 6. Control (saline water (1 ml/kg); IL-6 (ng/mL). *p< 0.05 when compared with control saline water group. ^#^p< 0.05 when compared with control DOX group.

### 3.6 Liver and Renal and Liver Function Biomarkers

Biochemical activities are represented in Table 1. In the liver, DOX alone increased AST by 35.4% and ALP by 46.7% compared to the control (normal) group. Co-treatment with UDHM12 + DOX reduced ALT by 29.2% relative to DOX-only. Co-treatment with NFD + DOX significantly lowered AST and ALT by 29.2% and 37.2%, respectively, compared to DOX alone. Also, renal function markers showed DOX elevated plasma creatinine, urea, and bilirubin by 118.7%, 56.4%, and 68.8%, respectively, compared to control. Co-treatments with UDHM12 + DOX reduced creatinine, urea, and bilirubin by 31.1%, 33%, and 29.8%, respectively, relative to DOX. NFD + DOX also lowered these markers by 38.3%, 34.9%, and 25.7%, respectively, compared to DOX only. In the plasma, DOX decreased albumin (ALB) and total protein (TP) by 32% compared to control while co-treatment with UDHM12 + DOX increased ALB and TP by 45.6% and 41.9%, respectively, relative to DOX alone. NFD + DOX increased ALB and TP by 49.4% and 44.5%, respectively, compared to DOX only. More so, DOX treatment alone did not significantly change lipid levels compared to control. However, co-treatments led to reductions in lipid markers in LDHM12 + DOX decreased TC by 86.1%, TG by 57.3%, and BIL by 26.1%, UDHM12 + DOX decreased TC by 52.9%, TG by 60.9%, and BIL by 55.4% and NFD + DOZ decreased TC by 46.8%, TG by 57.3%, and BIL by 20.7%.

**Table 1:** Effect of HM12 on biochemical parameters, oxidative and antioxidants parameters in normal and doxorubicin-treated male rats.

| Parameter | CONTROL | DOX | LDHM12 | UDHM12 | NFD | NFD + DOX | LDHM12 + DOX | UDHM12 + DOX |
| --- | --- | --- | --- | --- | --- | --- | --- | --- |
| AST | 16.61±0.64 | 22.49±1.09* | 17.10±0.80 | 17.96±0.53 | 12.92±1.09 | 13.92±0.27<sup>a</sup> | 19.01±0.35 | 16.81±0.39 |
| ALT | 40.67±9.04 | 46.20±0.98 | 40.10±2.89 | 41.37±2.86 | 40.30±2.42 | 33.60±0.29 | 38.55±0.49 | 35.09±0.72 |
| ALP | 14.51±1.48 | 21.29±2.62* | 15.62±1.72 | 16.03±2.17 | 16.32±2.93 | 13.38±0.78<sup>a</sup> | 15.28±1.35 | 15.07±0.21<sup>a</sup> |
| CREA | 40.07±5.68 | 86.70±11.2* | 51.97±4.12 | 42.10±0.92 | 45.27±1.47 | 53.50±4.63 | 61.85±5.45 | 59.75±2.34 |
| UREA | 6.97±0.38 | 10.90±0.12* | 6.87±0.43 | 6.80±0.35 | 6.20±0.06 | 7.10±0.06 | 7.70±0.35 | 7.30±0.29 |
| BILI | 1.73±0.44 | 2.92±0.33* | 2.37±0.15 | 2.45±0.09 | 2.47±0.09 | 2.17±0.03 | 2.45±0.03 | 2.05±0.32 |
| TC | 2.09±0.06 | 3.89±0.05* | 2.08±0.04 | 2.03±0.09 | 2.00±0.03 | 2.07±0.02<sup>a</sup> | 1.83±0.07<sup>a</sup> | 1.71±0.04<sup>b</sup> |
| TG | 0.42±0.09 | 0.82±0.06** | 0.49±0.08 | 0.40±0.09 | 0.43±0.03 | 0.35±0.03<sup>a</sup> | 0.36±0.05<sup>a</sup> | 0.32±0.06<sup>a</sup> |
| HDL | 1.63±0.02 | 0.92±0.04 | 1.08±0.01 | 1.33±0.02 | 1.01±0.01 | 1.11±0.02<sup>a</sup> | 1.16±0.02<sup>a</sup> | 1.43±0.04<sup>b</sup> |
| ALB | 39.90±1.39 | 27.10±1.09* | 34.47±0.43 | 41.27±1.65 | 38.40±0.35 | 41.87±0.55<sup>a</sup> | 30.65±1.99 | 39.45±0.95 |
| TP | 76.80±2.54 | 52.70±1.79 | 80.10±0.58 | 78.37±0.09 | 75.77±1.53 | 76.17±0.66<sup>a</sup> | 65.50±1.09 | 78.75±0.96<sup>a</sup> |
| MDAk | 1.42 ± 0.04 | 3.05 ± 0.03* | 1.93 ± 0.02 | 2.03 ± 0.03 | 2.07 ± 0.02 | 2.28 ± 0.04<sup>a</sup> | 2.33 ± 0.04 | 2.04 ± 0.02<sup>a</sup> |
| GSHk | 2.44 ± 0.03 | 1.11 ± 0.02* | 2.41 ± 0.03 | 2.77 ± 0.02 | 2.69 ± 0.03 | 2.28 ± 0.04<sup>a</sup> | 2.18 ± 0.04 | 2.72 ± 0.02<sup>b</sup> |
| CATk | 1.38 ± 0.03 | 0.84 ± 0.03** | 1.17 ± 0.02 | 1.81 ± 0.03 | 1.88 ± 0.02 | 1.58 ± 0.02<sup>a</sup> | 1.02 ± 0.02 | 1.57 ± 0.04<sup>a</sup> |
| SODk | 2.31 ± 0.05 | 1.07 ± 0.04* | 1.71 ± 0.04 | 2.70 ± 0.03 | 2.65 ± 0.03 | 1.82 ± 0.03<sup>a</sup> | 1.34 ± 0.03 | 1.71 ± 0.05<sup>a</sup> |
| MDA<sub>L</sub> | 1.31 ± 0.07 | 3.22 ± 0.05* | 2.11 ± 0.04 | 1.84 ± 0.04 | 1.69 ± 0.03 | 2.08 ± 0.04<sup>a</sup> | 3.07 ± 0.03 | 2.35 ± 0.05<sup>a</sup> |
| GSH<sub>L</sub> | 2.44 ± 0.04 | 1.29 ± 0.03* | 2.18 ± 0.03 | 2.55 ± 0.03 | 2.67 ± 0.04 | 1.53 ± 0.04 | 1.32 ± 0.06 | 1.59 ± 0.06 |
| CAT<sub>L</sub> | 0.86 ± 0.02 | 0.43 ± 0.02* | 0.92 ± 0.03 | 1.07 ± 0.02 | 1.31 ± 0.03 | 0.80 ± 0.03<sup>a</sup> | 0.51 ± 0.03 | 0.77 ± 0.02<sup>a</sup> |
| SOD<sub>L</sub> | 1.31 ± 0.03 | 1.02 ± 0.03* | 1.11 ± 0.02 | 1.39 ± 0.02 | 1.34 ± 0.04 | 1.41 ± 0.03 | 1.08 ± 0.03 | 1.34 ± 0.03 |

Results represented as Mean ± Standard Error of the Mean (SEM), with *N* = 6 animals per group.
Control (saline) group (10 ml/kg); i.p.: intraperitoneal; p.o. Per os (orall).
\**p* < 0.05 or \*\**p* < 0.001 when compared with control distilled water group. <sup>a</sup>*p* < 0.05 or <sup>b</sup>*p* < 0.001 when compared with DOX group.
AST: Aspartate Aminotransferase; ALT: Alanine Aminotransferase; ALP: Alkaline Phosphatase; BILI: Total Bilirubin; TC: Total Cholesterol; TG: Triglycerides; HDL: High Density Lipoprotein; ALB; Albumin; TP: Total Protein and Urea (mg/dL). MDA<sub>k/L</sub>: malondialdehyde (μmol/ml), GSH<sub>k/L</sub>: Kidney reduced glutathione (μmol/ml), SOD<sub>k/L</sub>: Superoxide dismutase (μmol/ml/min/mg protein), CAT<sub>k/L</sub>: Catalase (μmol/ml/min/mg protein).

### 3.7 Hepatic and Renal Lipid Peroxidation and Antioxidant Enzymes

Administration of DOX significantly elevated lipid peroxidation (LPO) levels in renal and hepatic tissues over 70% (Table 1). However, co-treatment with UDHM12 + DOX and NFD + DOX significantly reduced hepatic LPO levels by 27.3% and 35.4%, respectively, compared to DOX-only treatment. However, neither UDHM12 nor NFD was effective in mitigating DOX-induced renal MDA in the treated rats. DOX intoxication also resulted in a decrease in glutathione (GSH) levels: renal GSH was lowered by 54.5% and 47.1%, while hepatic GSH decreased by 23.3% and 18.6% (Table 1). Furthermore, DOX-only treatment reduced renal catalase (CAT) and superoxide dismutase (SOD) activities by 39.1% and 53.7%, respectively. Treatment with UDHM12 + DOX or NFD + DOX reversed these reductions, increasing renal CAT and SOD by more than half. Similarly, hepatic CAT and SOD activities decreased by 50% and 22.1% under DOX treatment compared to the normal saline control group. These decreases were ameliorated in the UDHM12 + DOX group (79.1% and 31.4%) and the NFD + DOX group (86% and 38.2%) relative to the DOX-only group.

### 3.8 Interactions of DOX, NFD and HM12 with human TOP2B (3QX3)

The binding affinity (kcal/mol) of doxorubicin, nifedipine and HM12 with Human TOP2β (3QX3) were 8.4, 6.4 and 7.1 at zero rmsd/lb and rmsd (Table 2). The protein-ligand interactions of DOX and chain A of 3QX3 showed conventional hydrogen at ALA 772, ARG 743 and TYR 846. Also, a Pi-Sigma was found at LUE and Alkyl or Pi-Alkyl was demonstrated at PHE 1019 and PRO 740 respectively. Several van der Waals interactions were seen at HIS 774, TYR 773, LYS 744, VAL 852, ASP 847, SER 771, LYS 739, SER 733, ASP 1020, GLY 1023, and TRP 856, GLU 855 as well as a carbon hydrogen bond at PHE 738. In the NFD and chain A of 3QX3, there were conventional hydrogen (ARG 743) and alkyl bonds at LEU 845 or PRO 740. Also, NFD-3QX3 interactions produced a Pi-Sigma bond at PHE 1019. Other notables interactions resulting from NFD-3QX3 were carbon hydrogen bond (GLY 1023) and van der Waals (GLU 855, TRP 856, SER 733, GLU 728 and GLU 853). More so, the interactions between H12 and 3QX3 led to Pi-alkyl or alkyl bond (ARG 729 and ILE 872) and conventional hydrogen bond at LYS 739 respectively. In addition, van der Waals results into TYR 773, SER 730, GLY 871, GLU 870, ALA 779, MET 782, ASN 786, THR 783, GLN 742, and GYL 741 (Supplementary Figure 2).

**Table 2.** HM12, Doxorubicin, nifedipine and their Binding Affinities with human topoisomerase II beta in DNA complex.

| Compound | Smile | Molecular<br>Formular | Molecular<br>Weight<br>(g/mol) | PubChem<br>CID | Binding<br>Affinity<br>(kcal/mol) | rms<br>d/lb | rmsd/<br>lb |
| --- | --- | --- | --- | --- | --- | --- | --- |
| Doxorubicin | C[C@H]1[C@H]([C@H](C[C@@H](O1)O[C@H]2C[C@@](CC3=C2C(=C4C(=C3O)C(=O)C5=C(C4=O)C(=CC=C5)OC)O)(C(=O)CO)O)N)O | C<sub>27</sub>H<sub>29</sub>NO | 543.5 | 31703 | 8.4 | 0 | 0 |
| Nifedipine | CC1=C(C(C(=C(N1)C)C(=O)OC)C2=CC=CC=C2[N+](=O)[O-])C(=O)OC | C<sub>17</sub>H<sub>18</sub>N<sub>2</sub>O<sub>6</sub> | 346.3 | 4485 | 6.4 | 0 | 0 |
| HM12 | CC(C(=O)OCCN1C(=C(C(C=2C(C(CCC12)(C)C)=O)C1=C(C(=CC(=C1)Br)Br)O)C(=O)[O-])C)=C | C<sub>25</sub>H<sub>26</sub>Br<sub>2</sub>NO<sub>6</sub> | 596.3 | - | 7.1 | 0 | 0 |

Binding Affinity n = 3

### 3.9 Histology

Histological sections of heart tissue from DOX-intoxicated rats revealed interlacing fascicles of cardiac myocytes (Figure 9). In several areas of the myocardium and associated blood vessels, there were characteristics inflammatory aggregates around red blood cells and indicative of severe vascular congestion and edema in the DOX-untreated group. Also, co-administration of LDHM12 + DOX showed interlacing fascicles of myocardial cells accompanied by inflammatory infiltrations and red blood cell accumulation. Conversely, no visible lesions were observed in rats treated with UDHM12 + DOX, or NFD + DOX when compared to the control DOX group, indicating potential protective effect this compound. More so, in the liver, DOX induced edema and prominent red blood cell aggregates, which were ameliorated by UDHM12 and NFD treatments (Figure 10). Further, HM12 or NFD did not significantly mitigate red blood cell aggregation or mild vascular congestion in the kidneys of DOX-treated rats (Figure 11), suggesting limited renoprotective efficacy.

**Figure 9.**
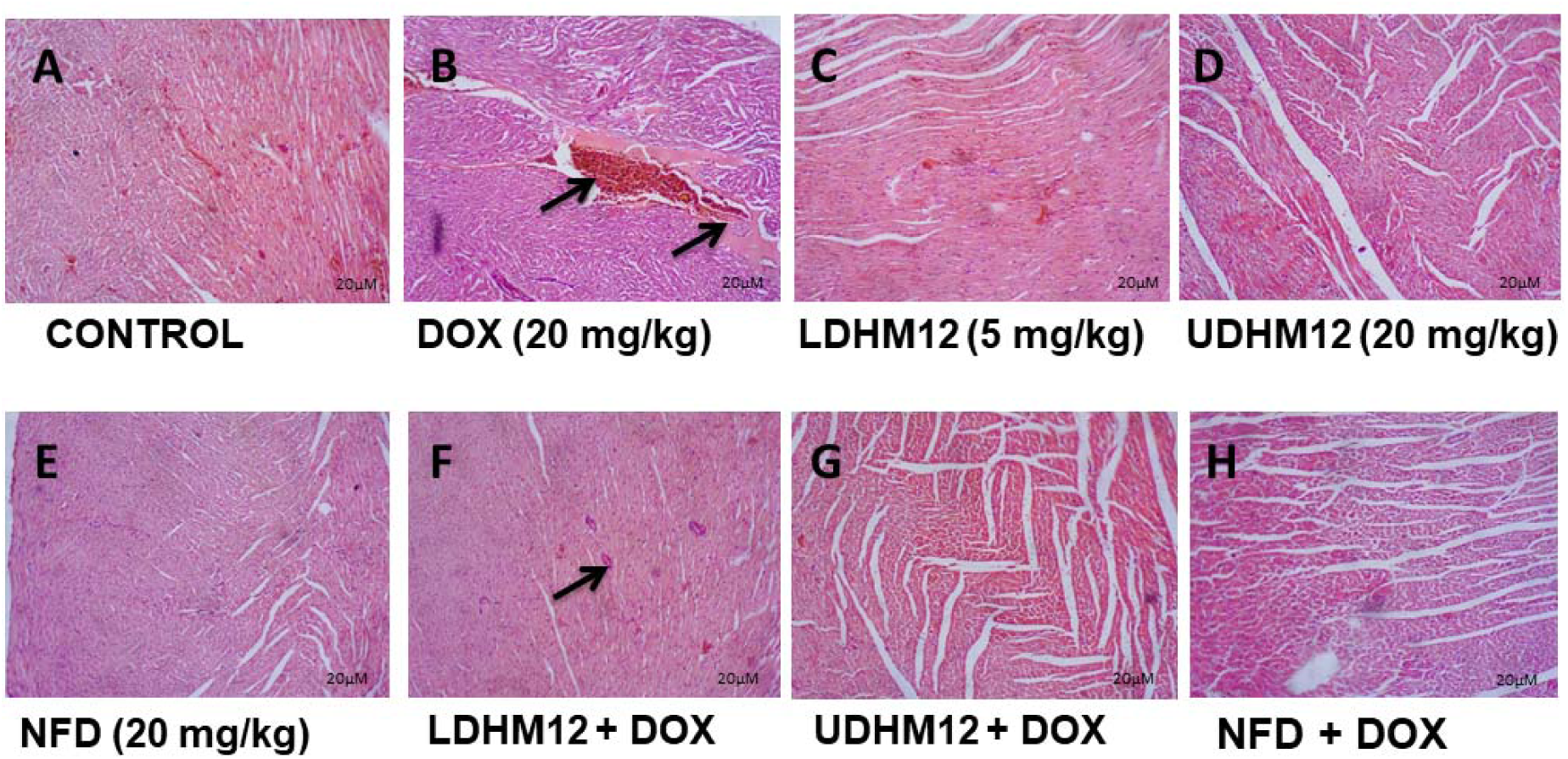
Histologic sections of rat heart muscle show interlacing fascicles of cardiac myocytes/ myocardial cells. No abnormalities are seen (A, C, D, E, G and H). Histologic sections of heart muscle show interlacing fascicles of cardiac myocytes/ myocardial cells with areas of the myocardium and vesicles showing several aggregates of red blood cells and smooth to slightly floccular pink fluid material common with severe vascular congestion with edema (B). LDHM12 + DOX, show interlacing fascicles of cardiac myocytes/ myocardial cells with extensive inflammation caused by aggregates of red blood cell myocardial inflammation (F). N: Nuclei; ID: Intercalated Disc; MF: Muscle Fibre; CV: Cytoplasmic Vacuole (H & E stain, ×100).

**Figure 10.**
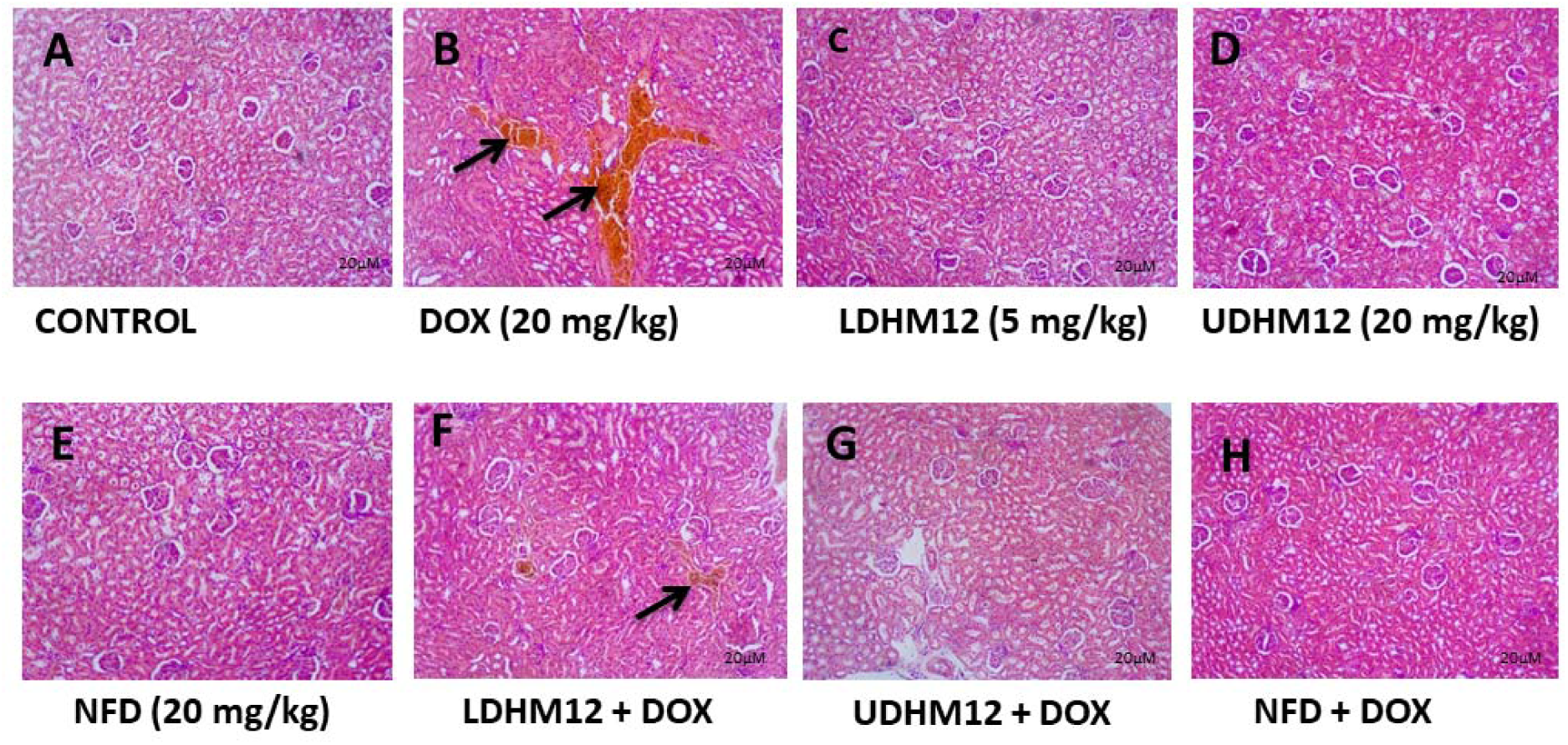
Histologic sections of liver tissue show parallel radially arranged plates of hepatocytes with Central vein (CV), portal vein (PV) and the basophilic portion with nucleus and the acidophilic cytoplasm of the acinar cells with no abnormalities are seen in A, C, D E, G and H. But, B, and F showed parallel radially arranged plates of hepatocytes, with the portal space and periportal zone filled with a smooth to slightly floccular pink fluid material common with edema and congested aggregates of red blood cells also seen. N: Nuclei; ID: Intercalated Disc; MF: Muscle Fibre; CV: Cytoplasmic Vacuole (H & E stain, ×100).

**Figure 11.**
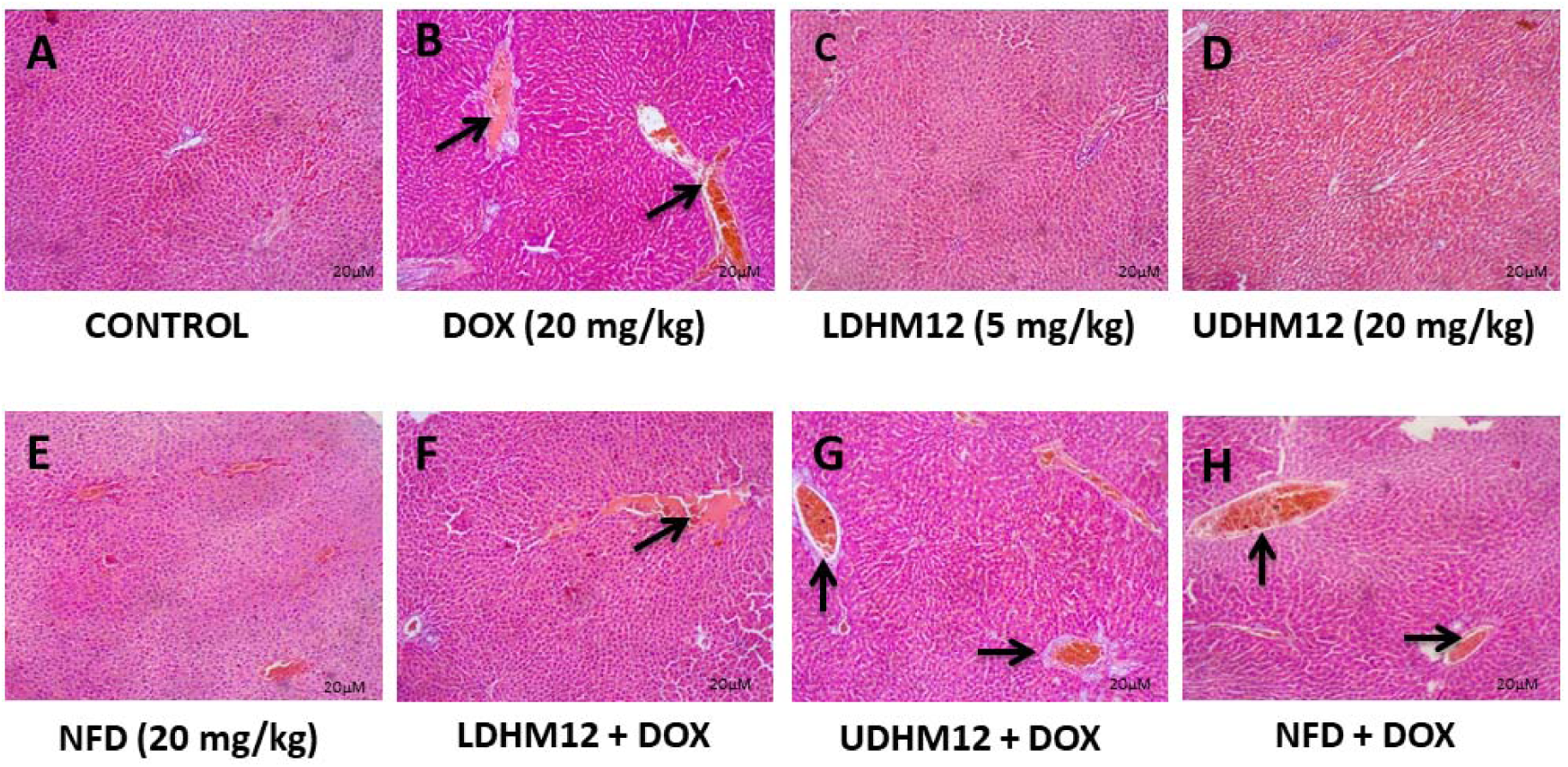
Histologic sections of well-defined kidney tissue show normocellular glomerular tufts disposed on a background containing renal tubules with no abnormalities are seen in A, C, D, and E. But, B sections showed vessels are filled with aggregates of red blood cells congesting vascular tissues. In F, G, and H His vessels are filled with aggregates of red blood cells with mild vascular congestion (H & E stain ×100).

## 4 DISCUSSION

DOX remains a cornerstone in the treatment of various malignancies. However, its clinical utility is significantly hindered by its cardiotoxic potential (Saleh et al., 2021). Chronic cardiomyopathy, a frequent adverse outcome in patients receiving DOX, limits its long-term therapeutic use (Rawat et al., 2021). This study investigates the cardioprotective potential of HM12, a 1,4-dihydropyridine (DHP) derivative with L-/T-type calcium channel-blocking properties, in mitigating DOX-induced cardiotoxicity in rats. Also, several mechanisms have been proposed for DOX-induced cardiomyopathy, including apoptosis triggered by oxidative radicals, membrane destabilization, mitochondrial dysfunction, and bioenergetic failure (Ralph et al., 2010; Antonucci et al., 2021; Sheibani et al., 2022). Lipid peroxidation and calcium homeostasis dysregulation are especially prominent among these mechanisms and are often coexistent in DOX-induced cardiac injury (Zhang et al., 2021). Genetic alterations, histaminergic activity, catecholaminergic surges, and immune dysregulation have also been implicated (Shepherd, 2003; Asnani, 2021; Yuan et al., 2021).

Despite early increases in mortality associated with DOX, early intervention has shown promise in improving outcomes (Gil-Gil et al., 2021). Consequently, various pharmacologic agents and natural compounds have been explored for their cardioprotective potential (Demerdash et al., 2021; Panneerpandian et al., 2021; Maleki et al., 2022). Antioxidants, such as vitamin E, N-acetylcysteine, coenzyme Q10, melatonin, and cysteamine, have shown varying degrees of effectiveness against DOX-induced toxicity (Quiles et al., 2002; Najafi et al., 2020; Abushouk et al., 2020).

DHPs including carvedilol, nifedipine, amlodipine, and felodipine, possess both vasodilatory and antioxidant properties that may mitigate DOX-induced cardiotoxicity (Hassan et al., 2021; Panneerpandian et al., 2021). In this study, we observed an increase in heart-to-body weight ratio in DOX-treated rats, consistent with prior evidence linking calcium dysregulation and cardiomyopathy (Awad et al., 2021; Shinlapawittayatorn et al., 2022). Treatment with high-dose HM12 and NFD moderately reduced heart weight, suggesting a protective effect, albeit limited. HM12’s potential modulation of intracellular calcium and oxidative stress pathways is under current investigation. DOX administration elevated plasma C-RP and LDH, markers of systemic inflammation and tissue damage. However, HM12 and NFD treatments significantly attenuated these elevations. DOX anticancer activity triggers oxidative stress and may impair intracellular mitochondrial actions resulting in ROS and subsequent impairment of redox homeostasis has been reported in vital organs including heart, kidneys and liver. Oxidative stress markers revealed increased MDA and decreased levels of GSH, CAT, and SOD in DOX-intoxicated rats, indicating severe oxidative injury (Kong et al., 2022; Zhang et al., 2021; Saleh et al., 2022). HM12 and NFD administration lowered MDA and improved antioxidant enzyme activity, further supporting HM12’s antioxidative action. Furthermore, DOX significantly elevated pro-inflammatory cytokines such as TNF-α and IL-6. While HM12 did not restore TNF-α levels, it significantly reduced IL-6, suggesting selective anti-inflammatory effects. Given that GSH is crucial for myocardial contraction, its depletion may exacerbate DOX-induced lethality (Saleh et al., 2022). From our results, HM12 at upper doses restored GSH and CAT, aligning with previous reports that DHPs exert cardioprotection through antioxidant and possibly β-adrenergic blockade or mitochondrial preservation (Mordente et al., 2012; Panneerpandian et al., 2021; Morciano et al., 2022). Although HM12 reduced lipid peroxidation and improved antioxidant status, it offer partial protection to the liver but did not reverse DOX-induced renal damage. DHP class is well known vasodilators with effects enhancing blood vessels. By this effort, studies on CCBs agents show considerable decrease in the liver calcium accumulation and demonstrate prevention cell injury with reversal in the centrilobular necrosis associated with hepatic toxicants have been documented (Scalzo et al., 2022). This highlights the complexity of organ-specific responses and suggests that HM12’s cytoprotective effects may be more cardiac-specific. While a modest reduction in alkaline phosphatase was observed, its clinical relevance remains unclear and warrants further investigation (Akindele et al., 2018).

In silico, NFD shared interactions similar Pi-Alkyl with DOX at PHE 1019 and PRO 740, and a conventional hydrogen bonding ARG 743 when interacted with human 3QX3 (Supplementary Figure 2). Also both DOX and NFD showed van der Waals interactions at SER 733, TRP 856, GLU 855. However, DOX van der Waals interaction at GLY 1023 was converted by NFD into a carbon hydrogen bond for the same. Both DOX and HM12 showed van der Waals at TYP 773.

However, HM12 had conventional hydrogen bond at LYS 739 in replacement to van der Waals interactions seen at TYR 773 in DOX interaction with human 3QX3. Additionally, HM12 interactions with human 3QX3 led to Pi-alkyl or alkyl bond (ARG 729 and ILE 872) and van der Waals results into SER 730, GLY 871, GLU 870, ALA 779, MET 782, ASN 786, THR 783, GLN 742, and GYL 741 (Supplementary Figure 2). Our results, both in vivo and in silico, align with previous reports indicating that DHP CCBs modulates DOX-induced cardiotoxicity (Santostasi et al., 1991; Belger et al., 2024;), potentially through overlapping protein-ligand interactions with human 3QX3 and the promotion of oxidative damage (Ikeda et al., 2019; Belger et al., 2024; Sripusanapan et al., 2025). Notably, mixed L- and T-type HM12 exhibited greater binding affinity than NFD and engaged in distinct interaction patterns with 3QX3 that were not observed with DOX which showed potential chemoprotective effects through mitigation of calcium-mediated damage. Further, it appears both potential vasodilation effects plus inhibition of T-type voltage gated calcium channel.

5 CONCLUSIONS

HM12 demonstrated cardioprotection potentially against DOX-induced toxicity by modulating oxidative stress, calcium dysregulation, and inflammation. It also influenced the hemodynamic activities of the hepato-renal axis. However, its limited ability to fully prevent multi-organ damage highlights the need for combination therapies or novel analogs with broader protective profiles. Continued investigation into the molecular mechanisms of HM12 will be crucial for its potential clinical application in reducing DOX-induced cardiotoxicity and improving the therapeutic index of this essential chemotherapeutic agent.

## Author contributions

KOE, GMG, OIO, AO, and EM: Investigation, Formal analysis, Writing - original draft, Writing - review and editing. KOE, GMG, OIO: Investigation. KOE, GMG, and AO: Conceptualization, Supervision. KOE, GMG, OIO: Conceptualization, Resources. KOE, GMG, OIO, AO, and EM- Conceptualization, Supervision, Validation, Project administration, Writing, Review and Editing. All authors contributed to the article and approved the submitted version.

## Ethics Statement

Ethical approval was obtained from University Health Research Ethics Committee (BUHREC 376/18). The authors gave their explicit consent to publish the manuscript. Oluwafemi Kale is available to contact any of the people involved in the research for further enquiry and information.

## Data Availability Statement

The data that support the findings of this study are available from the corresponding author upon reasonable request. Data will be made available on request.

## REFERENCES

Abushouk, A. I., Ismail, A., Salem, A. M. A., Afifi, A. M., & Abdel-Daim, M. M. (2017). Cardioprotective mechanisms of phytochemicals against doxorubicin-induced cardiotoxicity. Biomedicine & Pharmacotherapy, 90, 935–946.

Akindele, A. J., Oludadepo, G. O., Amagon, K. I., Singh, D., & Osiagwu, D. D. (2018). Protective effect of carvedilol alone and coadministered with diltiazem and prednisolone on doxorubicin and 5 fluorouracil induced hepatotoxicity and nephrotoxicity in rats. Pharmacology research & perspectives, 6(1), e00381.

Aloss, K., & Hamar, P. (2023). Recent Preclinical and Clinical Progress in Liposomal Doxorubicin. Pharmaceutics, 15(3), 893.

Antonucci, S., Di Lisa, F., & Kaludercic, N. (2021). Mitochondrial reactive oxygen species in physiology and disease. Cell Calcium, 94, 102344.

Asnani, A. (2021). Activating Autophagy to Prevent Doxorubicin Cardiomyopathy: The Timing Matters. Circulation research, 129(8), 801–803.

Awad, H. H., El-Derany, M. O., Mantawy, E. M., Michel, H. E., Mona, M., El-Din, R. A. S., … & El-Demerdash, E. (2021). Comparative study on beneficial effects of vitamins B and D in attenuating doxorubicin induced cardiotoxicity in rats: Emphasis on calcium homeostasis. Biomedicine & Pharmacotherapy, 140, 111679.

Aygün Cevher, H., Schaller, D., Gandini, M. A., Kaplan, O., Gambeta, E., Zhang, F. X., … & Gündüz, M. G. (2019). Discovery of Michael acceptor containing 1, 4-dihydropyridines as first covalent inhibitors of L-/T-type calcium channels. Bioorganic Chemistry, 91, 103187.

Bertram-Ralph, E., & Amare, M. (2021). Pharmacological modulation of cardiac function and control of blood vessel calibre. Anaesthesia & Intensive Care Medicine, 22(11), 738–748.

Bhatt, K. S., Singh, A., Marwaha, G. S., Ravendranathan, N., Sandhu, I. S., Kim, K., … & Singh, K. K. (2025). Different Mechanisms in Doxorubicin-Induced Neurotoxicity: Impact of BRCA Mutations. International Journal of Molecular Sciences, 26(10), 4736.

Bisht, A., Avinash, D., Sahu, K. K., Patel, P., Das Gupta, G., & Kurmi, B. D. (2025). A comprehensive review on doxorubicin: mechanisms, toxicity, clinical trials, combination therapies and nanoformulations in breast cancer. Drug Delivery and Translational Research, 15(1), 102–133.

Gil-Gil, M. J., Bellet, M., Bergamino, M., Morales, S., Barnadas, A., Manso, L., … & Pernas, S. (2021). Long-Term Cardiac Safety and Survival Outcomes of Neoadjuvant Pegylated Liposomal Doxorubicin in Elderly Patients or Prone to Cardiotoxicity and Triple Negative Breast Cancer. Final Results of the Multicentre Phase II CAPRICE Study. Frontiers in oncology, 11.

Hassan, M. Q., Akhtar, M. S., Afzal, O., Hussain, I., Akhtar, M., Haque, S. E., & Najmi, A. K. (2020). Edaravone and benidipine protect myocardial damage by regulating mitochondrial stress, apoptosis signalling and cardiac biomarkers against doxorubicin-induced cardiotoxicity. Clinical and Experimental Hypertension, 42(5), 381–392.

Heravi, M. M., & Zadsirjan, V. (2022). Construction and Aromatization of Hantzsch 1, 4 Dihydropyridines under Microwave Irradiation: A Green Approach. ChemistrySelect, 7(3), e202104032.

Huber, K., Mestres Arenas, A., Fajas, L., & Leal Esteban, L. C. (2021). The multifaceted role of cell cycle regulators in the coordination of growth and metabolism. The FEBS Journal, 288(12), 3813–3833.

Iqbal M, Dubey K, Anwer T, Ashish A, Pillai K (2008) Protective effects of telmisartan against acute doxorubicin-induced cardiotoxicity in rats. Pharmacol Rep 60(3):382–390

Kale, O. E., Awodele, O., Ogundare, T. F., & Ekor, M. (2017). Amlodipine, an L type calcium channel blocker, protects against chlorpromazine induced neurobehavioural deficits in mice. Fundamental & clinical pharmacology, 31(3), 329–339.

Kamińska, K., & Cudnoch-Jędrzejewska, A. (2023). A review on the neurotoxic effects of doxorubicin. Neurotoxicity research, 41(5), 383–397.

Karthick, R., Velraj, G., Pachamuthu, M. P., & Karthikeyan, S. (2022). Synthesis, spectroscopic, DFT, and molecular docking studies on 1, 4-dihydropyridine derivative compounds: a combined experimental and theoretical study. Journal of Molecular Modeling, 28(1), 1–15.

Kilkenny, C., Browne, W. J., Cuthill, I. C., Emerson, M., & Altman, D. G. (2010). Improving bioscience research reporting: The ARRIVE guidelines for reporting animal research. PLoS Biology, 8(6), e1000412.

Kong, C. Y., Guo, Z., Song, P., Zhang, X., Yuan, Y. P., Teng, T., … & Tang, Q. Z. (2022). Underlying the Mechanisms of Doxorubicin-Induced Acute Cardiotoxicity: Oxidative Stress and Cell Death. International journal of biological sciences, 18(2), 760.

Le Cesne, A., Ouali, M., Leahy, M. G., Santoro, A., Hoekstra, H. J., Hohenberger, P., … & Van Der Graaf, W. T. A. (2014). Doxorubicin-based adjuvant chemotherapy in soft tissue sarcoma: pooled analysis of two STBSG-EORTC phase III clinical trials. Annals of Oncology, 25(12), 2425–2432.

Liu, L. L., Li, Q. X., Xia, L., Li, J., & Shao, L. (2007). Differential effects of dihydropyridine calcium antagonists on doxorubicin-induced nephrotoxicity in rats. Toxicology, 231(1), 81–90.

Maleki Dana, P., Sadoughi, F., Asemi, Z., & Yousefi, B. (2022). The role of polyphenols in overcoming cancer drug resistance: a comprehensive review. Cellular & Molecular Biology Letters, 27(1), 1–26.

Minotti, G., Menna, P., Salvatorelli, E., Cairo, G., & Gianni, L. (2004). Anthracyclines: molecular advances and pharmacologic developments in antitumor activity and cardiotoxicity. Pharmacological reviews, 56(2), 185–229.

Morciano, G., Rimessi, A., Patergnani, S., Vitto, V. A., Danese, A., Kahsay, A., … & Pinton, P. (2022). Calcium dysregulation in heart diseases: Targeting calcium channels to achieve a correct calcium homeostasis. Pharmacological Research, 177, 106119.

Mordente, A., Meucci, E., Silvestrini, A., Martorana, G. E., & Giardina, B. (2012). Anthracyclines and mitochondria. Advances in Mitochondrial Medicine, 385–419.

Najafi, Masoud, Mohammad Reza Hooshangi Shayesteh, Keywan Mortezaee, Bagher Farhood, and Hamed Haghi-Aminjan. “The role of melatonin on doxorubicin-induced cardiotoxicity: a systematic review.” Life sciences 241 (2020): 117173.

Panneerpandian, P., Rao, D. B., & Ganesan, K. (2021). Calcium channel blockers lercanidipine and amlodipine inhibit YY1/ERK/TGF-β mediated transcription and sensitize the gastric cancer cells to doxorubicin. Toxicology in Vitro, 74, 105152.

Quiles, J. L., Huertas, J. R., Battino, M., Mataix, J., & Ramırez-Tortosa, M. C. (2002). Antioxidant nutrients and adriamycin toxicity. Toxicology, 180(1), 79–95.

Ralph, S. J., Rodríguez-Enríquez, S., Neuzil, J., & Moreno-Sánchez, R. (2010). Bioenergetic pathways in tumor mitochondria as targets for cancer therapy and the importance of the ROS-induced apoptotic trigger. Molecular aspects of medicine, 31(1), 29–59.

Rawat, P. S., Jaiswal, A., Khurana, A., Bhatti, J. S., & Navik, U. (2021). Doxorubicin-induced cardiotoxicity: An update on the molecular mechanism and novel therapeutic strategies for effective management. Biomedicine & Pharmacotherapy, 139, 111708.

Saleh Ahmed, A. S. (2022). Potential protective effect of catechin on doxorubicin-induced cardiotoxicity in adult male albino rats. Toxicology Mechanisms and Methods, 32(2), 97–105.

Saleh, Y., Abdelkarim, O., Herzallah, K., & Abela, G. S. (2021). Anthracycline-induced cardiotoxicity: mechanisms of action, incidence, risk factors, prevention, and treatment. Heart failure reviews, 26(5), 1159–1173.

Sheibani, M., Azizi, Y., Shayan, M., Nezamoleslami, S., Eslami, F., Farjoo, M. H., & Dehpour, A. R. (2022). Doxorubicin-Induced Cardiotoxicity: An Overview on Pre-clinical Therapeutic Approaches. Cardiovascular Toxicology, 1-19.

Shepherd, G. M. (2003). Hypersensitivity reactions to chemotherapeutic drugs. Clinical reviews in allergy & immunology, 24(3), 253–262.

Shinlapawittayatorn, K., Chattipakorn, S. C., & Chattipakorn, N. (2022). The effects of doxorubicin on cardiac calcium homeostasis and contractile function. Journal of Cardiology.

Yuan, S. J., Wang, C., Xu, H. Z., Liu, Y., Zheng, M. Y., Li, K., … & Chen, X. (2021). Conjugation with nanodiamonds via hydrazone bond fundamentally alters intracellular distribution and activity of doxorubicin. International Journal of Pharmaceutics, 606, 120872.

Zhang, J., Duan, D., Song, Z. L., Liu, T., Hou, Y., & Fang, J. (2021). Small molecules regulating reactive oxygen species homeostasis for cancer therapy. Medicinal Research Reviews, 41(1), 342–394.

Zhang, J., Xiao, M., Wang, S., Guo, Y., Tang, Y., & Gu, J. (2021). Molecular mechanisms of doxorubicin-induced cardiotoxicity: Novel roles of sirtuin 1-mediated signaling pathways. Cellular and Molecular Life Sciences, 78(7), 3105–3125.

Scalzo, N., Canastar, M., & Lebovics, E. (2022). Part 1: disease of the heart and liver: a relationship that cuts both ways. Cardiology in Review, 30(3), 111–122.

Linders, A. N., Dias, I. B., López Fernández, T., Tocchetti, C. G., Bomer, N., & Van der Meer, P. (2024). A review of the pathophysiological mechanisms of doxorubicin-induced cardiotoxicity and aging. npj Aging, 10(1), 9

Bhutani, V., Varzideh, F., Wilson, S., Kansakar, U., Jankauskas, S. S., & Santulli, G. (2025). Doxorubicin-Induced Cardiotoxicity: A Comprehensive Update. Journal of Cardiovascular Development and Disease, 12(6), 207.

Vejpongsa, P., & Yeh, E. T. H. (2014). Topoisomerase 2β: a promising molecular target for primary prevention of anthracycline induced cardiotoxicity. Clinical Pharmacology & Therapeutics, 95(1), 45–52.

Jones, K. E., Hayden, S. L., Meyer, H. R., Sandoz, J. L., Arata, W. H., Dufrene, K., … & Kaye, A. D. (2024). The evolving role of calcium channel blockers in hypertension management: pharmacological and clinical considerations. Current Issues in Molecular Biology, 46(7), 6315–6327.

Delint-Ramirez, I., Konada, L., Heady, L., Rueda, R., Jacome, A. S. V., Marlin, E., … & Madabhushi, R. (2022). Calcineurin dephosphorylates topoisomerase IIβ and regulates the formation of neuronal-activity-induced DNA breaks. Molecular cell, 82(20), 3794–3809.

Ganapathi, R. N., & Ganapathi, M. K. (2013). Mechanisms regulating resistance to inhibitors of topoisomerase II. Frontiers in pharmacology, 4, 89.

Sripusanapan, A., Piriyakulthorn, C., Apaijai, N., Chattipakorn, S. C., & Chattipakorn, N. (2025). Ivabradine ameliorates doxorubicin-induced cardiotoxicity through improving mitochondrial function and cardiac calcium homeostasis. Biochemical Pharmacology, 236, 116881.

Ikeda, S., Matsushima, S., Okabe, K., Ikeda, M., Ishikita, A., Tadokoro, T., … & Tsutsui, H. (2019). Blockade of L-type Ca2+ channel attenuates doxorubicin-induced cardiomyopathy via suppression of CaMKII-NF-κB pathway. Scientific reports, 9(1), 9850.

Belger, C., Abrahams, C., Imamdin, A., & Lecour, S. (2024). Doxorubicin-induced cardiotoxicity and risk factors. IJC Heart & Vasculature, 50, 101332.

Santostasi, G., Kutty, R. K., & Krishna, G. (1991). Increased toxicity of anthracycline antibiotics induced by calcium entry blockers in cultured cardiomyocytes. Toxicology and applied pharmacology, 108(1), 140–149.

