## Supplementary Figure 1 for "A new 1,4-Dihydropyridine-Based L-/T-Type Calcium Channel Inhibitor, HM12, Provides in Vivo and in silico Cardioprotective Effects in Doxorubicin-Treated Rats": Summplementary-Figures.docx

(i)
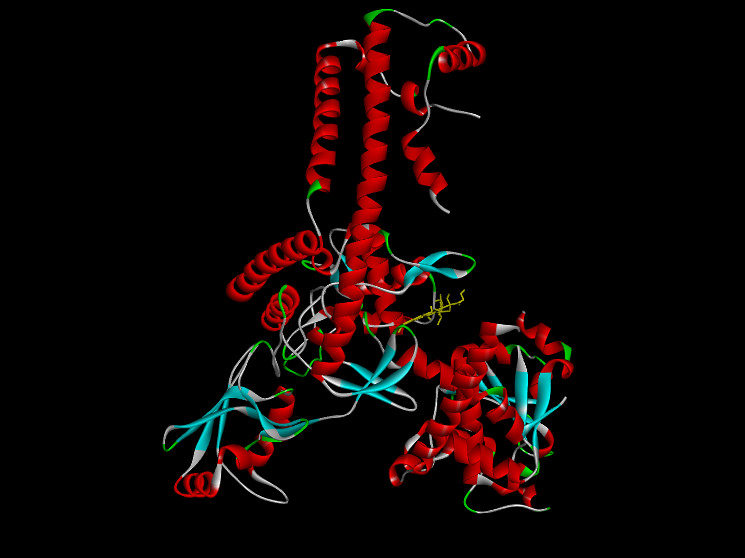
 (ii)
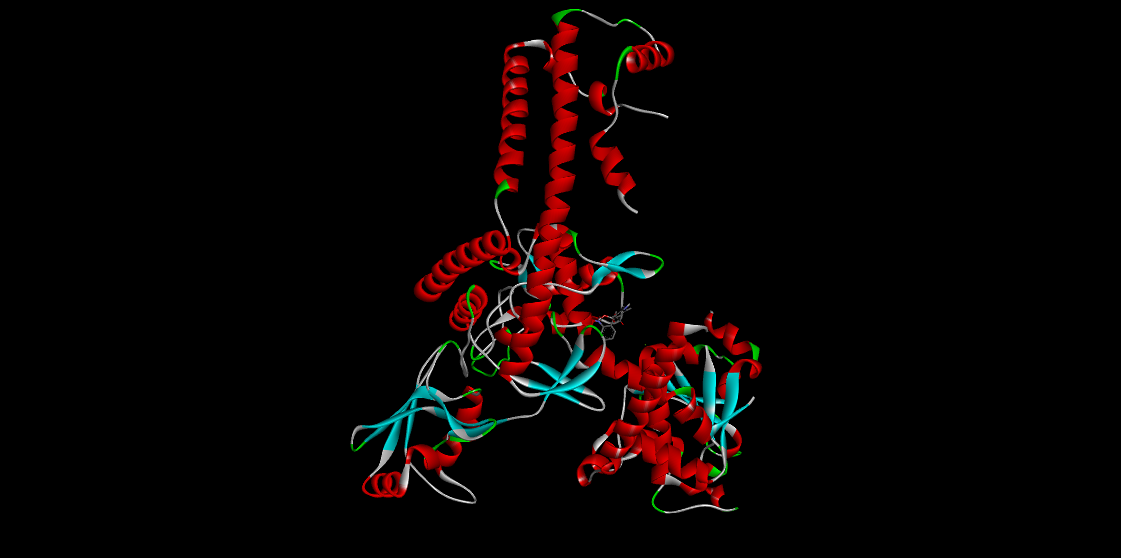
 (iii)
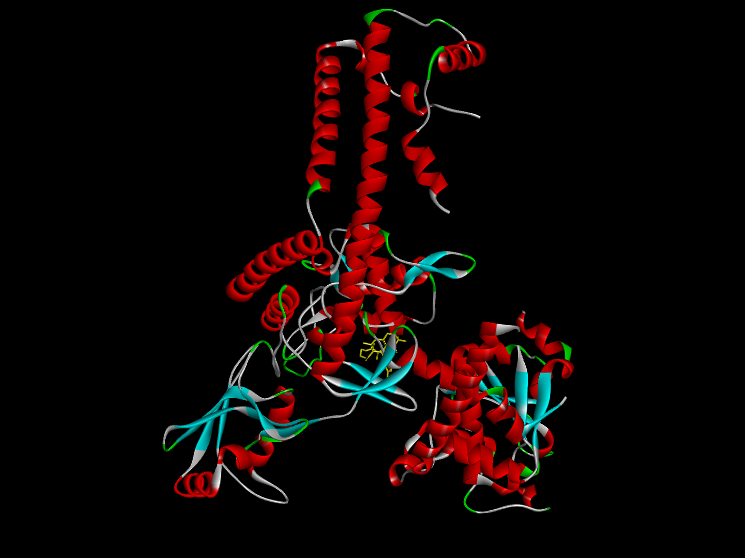

Supplementary Figure 1. 3D interaction of of purified chain A Human topoisomerase II beta in complex with DNA (3QX3) with (i) Doxorubicin (ii) Nifedipine and (iii) HM12

(i)
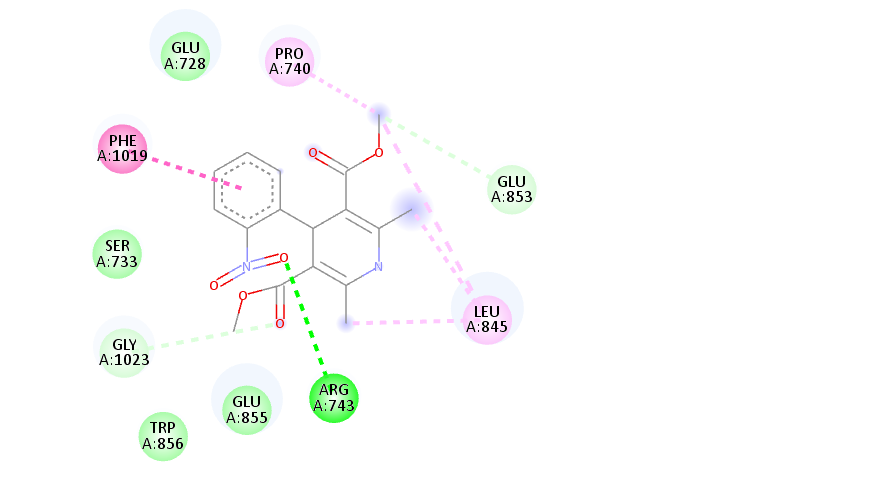

(ii)
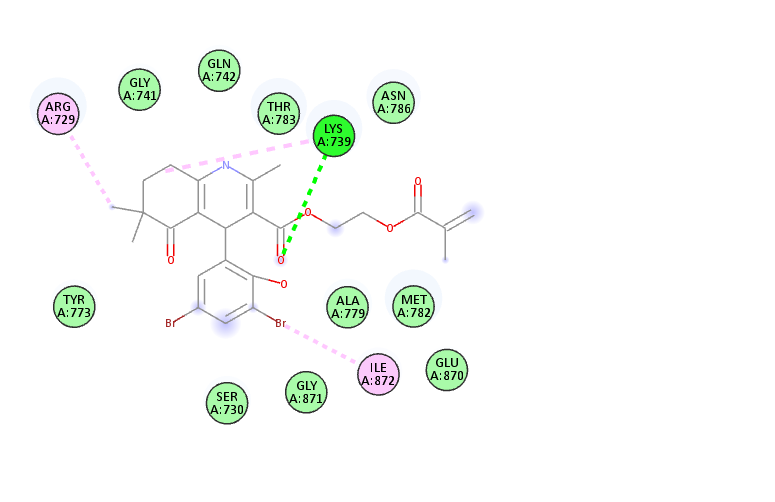

(iii)
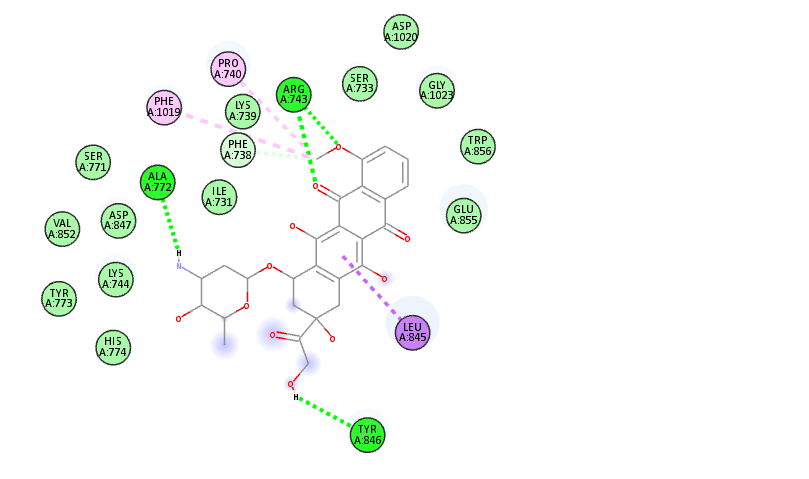

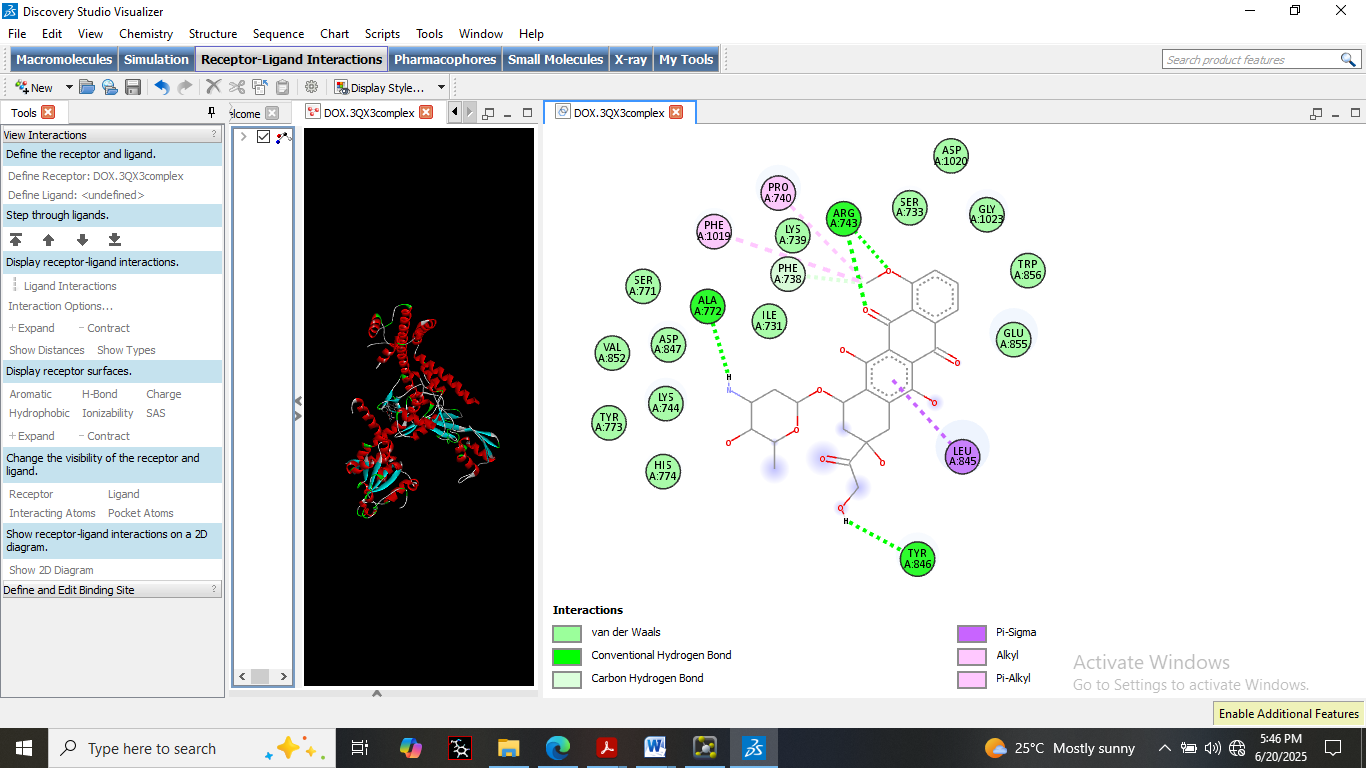

Supplementary Figure 2. 2D interaction of purified chain A of Human topoisomerase II beta in complex with DNA (3QX3) with (i) Doxorubicin (ii) Nifedipine and (iii) HM12
